# Five millennia of mitochondrial introgression in Atlantic bluefin tuna identified using ancient DNA

**DOI:** 10.1101/2024.03.24.586294

**Authors:** Emma Falkeid Eriksen, Adam Jon Andrews, Svein Vatsvåg Nielsen, Per Persson, Estrella Malca, Vedat Onar, Veronica Aniceti, Gäel Piquès, Federica Piattoni, Francesco Fontani, Martin Wiech, Keno Ferter, Oliver Kersten, Giada Ferrari, Alessia Cariani, Fausto Tinti, Elisabetta Cilli, Lane M. Atmore, Bastiaan Star

## Abstract

Mitogenomic (MT) introgression between species is readily documented in marine fishes. Such introgression events may either be long-term natural phenomena or the result of human-driven shifts in spatial distributions of previously separated species. Determining the drivers behind MT introgression is stymied by the difficulty of directly observing patterns of interbreeding over long timescales. Using ancient DNA spanning five millennia, we here investigate the long-term presence of MT introgression from Pacific bluefin tuna (*Thunnus orientalis*) and albacore (*Thunnus alalunga*) into Atlantic bluefin tuna (*Thunnus thynnus*), a species with extensive exploitation history and observed shifts in abundance, and demographic distribution. Comparing ancient (n=130) and modern (n=78) mitogenomes of specimens covering most of the range of Atlantic bluefin tuna we detect no significant spatial or temporal population structure. This lack of spatiotemporal genomic differentiation is indicative of ongoing gene flow between populations and large effective population sizes over millennia. Moreover, we identify introgressed MT genomes in ancient specimens up to 5000 years old and find that this rate of introgression has remained similar through time. We therefore conclude that MT introgression in the Atlantic bluefin tuna is to date unaffected by anthropogenic impacts. By providing the oldest example of directly observed MT introgression in the marine environment, our results highlight the utility of ancient DNA to obtain temporal insights in the long-term persistence of such phenomena.

## Introduction

Genetic introgression is the integration of genetic material from one parent species into another following interspecific hybridization and backcrossing (Rhymer & Simberloff, 1996). Although frequency of the phenomenon across biological systems remains debated, the increased use of next-generation sequencing across non-model taxa has revealed introgression to be a more common phenomenon in nature than previously thought (Dagilis et al., 2022). The majority of documented introgression in animals involves introgression of the mitochondrial (MT) genome (Sloan et al., 2017; Pons et al., 2014; Toews & Brelsford, 2012). Typically inherited maternally in vertebrates, the non-recombining introgressed mitogenome remains largely intact over time (Seixas et al., 2018; Brown, 2008). The presence of introgressed MT haplotypes can cause significant bias when using mitogenomic data to describe a species demographic properties or evolutionary history. Even rare hybridization events can result in the presence of whole MT haplotypes that do not accurately reflect the typical history or demography of the taxon. For example, the presence of introgressed MT haplotypes may dominate genealogies with recent dispersal history and thereby overshadow genetic signals from past dispersal events (Sloan et al., 2017; Ballard & Whitlock, 2004). Presence of introgressed, heterospecific alleles and haplotypes will also affect population genomic analyses by inflating measures of genetic diversity and divergence (Oosting et al., 2023; Rodriguez & Krug, 2022; Wang et al., 2022; Hawks, 2017). Avoiding such inflation is important because these statistics can influence management choices (Willi et al., 2022; Hohenlohe et al., 2021; Kardos et al., 2021) and inflated measures of genetic diversity or effective population size may exaggerate the genetic robustness of a truly vulnerable population.

Marine fish hybridize according to their ecologies and life history strategies, thus the rate of introgression will vary according to migration behavior, spawning site overlap, fecundity, spawning ontology, and offspring survival (Montanari et al., 2016; Gardner, 1997; Hubbs, 1955). In the economically important redfish (*Sebastes spp.*), high rates of introgressive hybridization (15% of all samples) have been found between two species (*S. fasciatus* and *S. mentella*) that live sympatrically in hybrid zones and yet maintain their morphology, resembling one of the parent species (Benestan et al., 2021; Roques et al., 2001). Likewise, introgression has been observed in European seabass (*Dicentrarchus labrax*) (Duranton et al., 2020; Vandeputte et al., 2019), capelin (*Mallotus villosus*) (Cayuela et al., 2020; Colbeck et al., 2011), European anchovy (*Engraulis encrasicolus*) (Le Moan et al., 2016), Australasian snapper (*Chrysophrys auratus*) (Oosting et al., 2023) and Atlantic and Pacific herring (*Clupea harengus* and *C. pallasii*) (Semenova, 2020).

Formation of hybrid zones after recent range shifts induced by contemporary climate change have already been observed in a number of species (Kersten et al. 2023; Ottenburghs, 2021; Garroway et al., 2010; Taylor & Larson, 2019; Ryan et al., 2018) including marine fish (Muhlfeld et al., 2014; Potts et al., 2014). The formation of such hybrid zones can have both deleterious and advantageous effects. For instance, in trout, warmer freshwater temperatures and lower precipitation is expected to increase introgressive hybridization between native European brown trout (*Salmo trutta*) and released non-native brown trout in Mediterranean rivers, potentially leading to loss of local genetic variants (Vera et al., 2023). Yet in rainbowfish (*Melanotaenia* spp.), it has been suggested that introgressive hybridization contributes to climate change resiliency by incorporating potentially adaptive genetic variation (Brauer et al., 2023; Turbek & Taylor, 2023). Regardless of the evolutionary consequences, knowledge about the *timing* of the introgression is necessary to understand if it is anthropogenic impacts that increase rates of hybridization, thereby positively or negatively altering the adaptive potential of species (Xuereb et al., 2021; Hoffmann & Sgrò, 2011).

In this study, we investigate introgression in the ecologically and economically important Atlantic bluefin tuna (*Thunnus thynnus*, Linneaus 1758), a highly migratory marine predatory fish distributed across the Atlantic Ocean (SCRS, 2023; Block, 2019; Nøttestad et al., 2020). Atlantic bluefin exhibits strong natal homing behavior (Brophy et al., 2016; Boustany et al., 2008; Block et al., 2005) and is therefore managed as two separate stocks: the larger Eastern stock spawning predominantly in the Mediterranean, and a smaller Western stock spawning predominantly in the Gulf of Mexico (ICCAT, 2023). Recent studies, however, have demonstrated weak genetic divergence in Atlantic bluefin and the existence of a previously unknown spawning ground in the Slope Sea where the stocks seem to interbreed (Diaz-Arce et al., 2024; Aalto et al., 2023; Andrews et al., 2021; Rodríguez-Ezpeleta et al., 2019), thereby challenging the assumption of two reproductively isolated populations. After severe international overfishing of Atlantic bluefin in the last century, the Eastern Atlantic bluefin stock has at present efficiently recovered due to strict management measures and favorable oceanographic conditions in the recent decade (ICCAT 2022a, 2022b) followed by improved recruitment with a series of very strong year classes (Reglero et al. 2018; Garcia et al. 2013; ICCAT 2023). The Atlantic bluefin has an extensive exploitation history, starting in the early Neolithic (ca. 6000 BCE), expanding through the Greek and Roman era and developing into an intense fishing industry towards the end of the last millennium (Andrews et al., 2022). Consequently, Atlantic bluefin stocks were depleted by the 21st century (Block, 2019; MacKenzie et al., 2009), leading to shifts in its demographic distribution and foraging behavior (Di Natale, 2015; MacKenzie et al., 2014; Andrews et al., 2023a; Worm & Tittensor, 2011). These distributional changes, as well as the establishment of potentially new spawning grounds may affect the rate of introgression in the species by providing increased or decreased opportunities for hybridization.

MT introgression within the Thunnus genus has been well documented, and together with the lack of reliable phylogenetically informative markers to distinguish the species, contributed to an unresolved phylogeny within the genus until recently (Alvarado Bremer et al., 1997; Chow et al., 2006; Chow & Kishino, 1995; Díaz-Arce et al., 2016; Santini et al., 2013; Viñas & Tudela, 2009). The Atlantic bluefin was previously thought to be a subspecies of Northern bluefin tuna together with Pacific bluefin (*Thunnus orientalis,* Temminck and Schlegel 1844). The bluefins are now regarded as distinct species (Chow et al., 2006; Díaz-Arce et al., 2016) with non-overlapping ranges (Tseng et al., 2011) (see Figure S1). The Pacific bluefin, distributed across the North Pacific Ocean, is closely related to albacore tuna (*Thunnus alalunga*, Bonnaterre 1788) in mitochondrial phylogenies (Gong et al., 2017; Viñas & Tudela, 2009; Chow et al., 2006). Albacore tuna is found in both the Pacific, Indian and Atlantic Oceans, including the Mediterranean Sea, typically preferring warmer waters than the Pacific and Atlantic bluefins, but with largely overlapping ranges and spawning areas (Chow & Ushiama, 1995; Saber et al., 2015). In the most recent and resolved nuclear phylogeny, the albacore tuna occurs as the sister-species to the other Thunnus species, and the Pacific and Atlantic bluefin form a monophyletic group (Díaz-Arce et al., 2016; Ciezarek et al. 2018) (see Figure S1).

MT introgression has been demonstrated from both albacore and Pacific bluefin into the Atlantic bluefin and from the Atlantic bluefin and albacore into the Pacific bluefin, but no MT introgression has been found in albacore (Alvarado Bremer et al., 2005; Carlsson et al., 2004; Chow & Kishino, 1995; Rooker et al., 2007; Viñas et al., 2003, 2011; Viñas & Tudela, 2009; Chow et al., 2006; Chow & Inoue, 1993). In the Atlantic bluefin, the rates of introgression from either albacore or Pacific bluefin are similar at around 2-5% (Rooker et al., 2007; Viñas & Tudela, 2009). Nonetheless, it is unclear when these introgression events started to occur or whether they are ongoing. In addition to the distributional shifts likely caused by high fishing pressures, it is possible that climate warming has contributed to novel opportunities for introgression in recent decades. The distribution of Atlantic bluefin over the last century has fluctuated with temperature (Faillettaz et al., 2019; Ravier & Fromentin, 2004), and ocean warming has been implicated in altering migration patterns, spawning ontology, and habitats of the Atlantic bluefin (Diaz-Arce et al., 2023; Faillettaz et al., 2019; Fiksen & Reglero, 2022; Muhling et al., 2011). Determining the timing of hybridization events can therefore shed light on the drivers of introgression.

DNA extracted from historical or ancient samples (hDNA or aDNA) allows us to directly investigate the timing of introgression events and to elucidate potential changes in population structure and genetic diversity over time (Kersten et al. 2023). Fish bones have physiological qualities that may increase the likelihood of finding well preserved DNA (Ferrari et al., 2021; Kontopoulos et al., 2019; Szpak, 2011) allowing for whole genome sequencing (Star et al., 2017), even from very limited amounts of bone (e.g. <10 mg) (Atmore et al., 2023). Here we use such aDNA methods to analyze mitogenomes from 130 ancient and 78 modern Atlantic bluefin spanning a period of approximately 5000 years (Figure 1). By sampling both before and after the period of heavy exploitation (1970-2007) and ongoing climate change, we investigate spatiotemporal patterns of genetic diversity and provide the longest historical time series of the presence of MT introgression in the marine environment to date.

**Figure 1:**
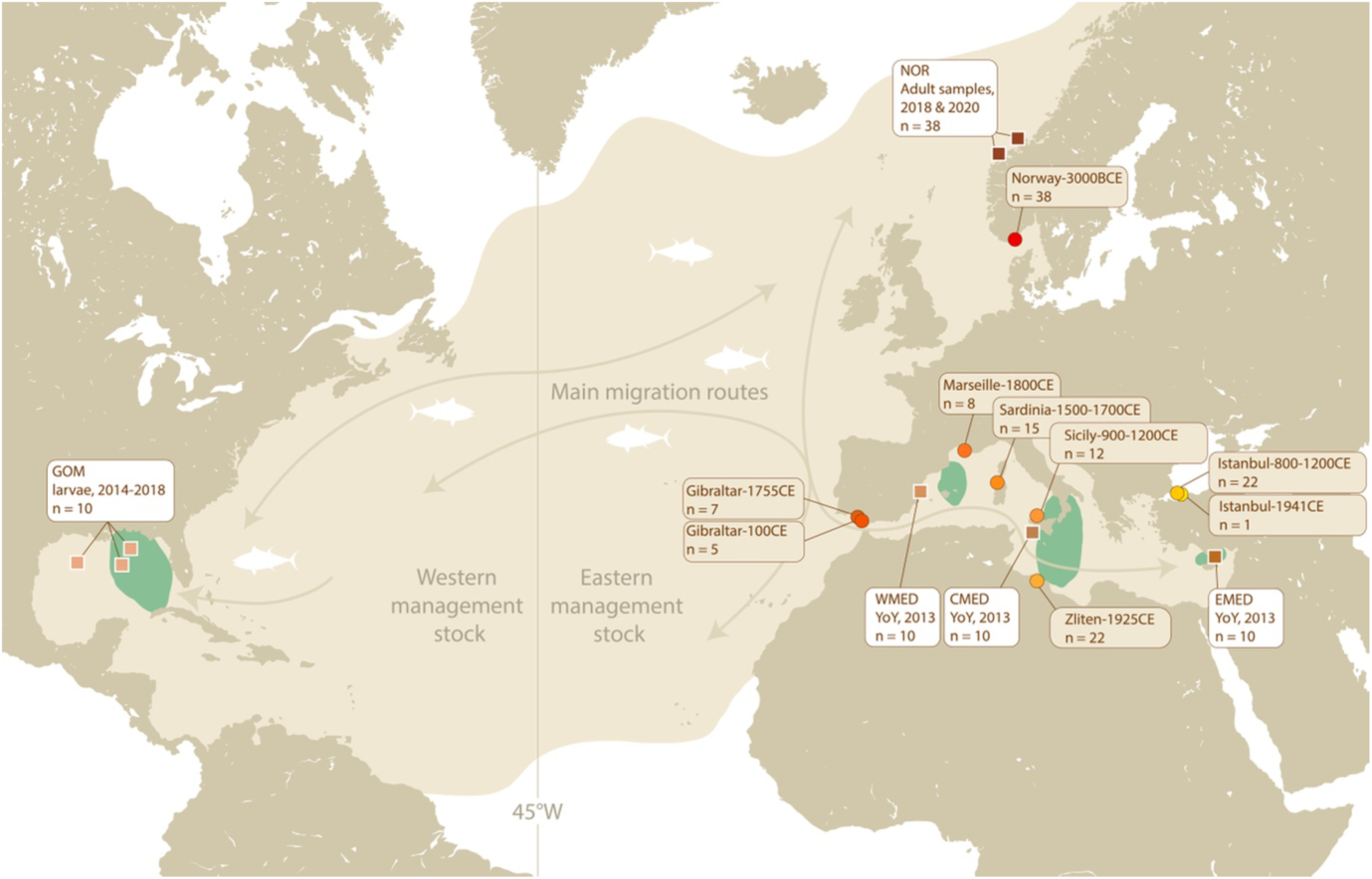
Distribution of the Atlantic bluefin tuna, including spawning areas (green) currently considered by management (adapted from IMR (2021)). The equal-distance line (45°W) separates the Eastern and Western stocks for management purposes. Sample locations of modern (squares, white boxes) and ancient tuna (circles, brown boxes) used in this study are indicated on the map. Arrows indicate the main migration routes of adult Atlantic bluefin (adapted from Fromentin et al., 2014). GOM = Gulf of Mexico, NOR = Norway, WMED = Western Mediterranean, CMED = Central Mediterranean, EMED = Eastern Mediterranean. YoY = young-of-the-year.

## Methods

### Collection, extraction, and sequencing of ancient samples from Norway

38 Neolithic (ca. 3000 BCE) tuna bones from the south of Norway were obtained from three archaeological excavations at Jortveit from 2018 to 2020. Bones were found at varying depths (42-130 cm) in six of nine total trenches and were estimated to be from 3700-2500 BCE based on radiocarbon dating of wood and charcoal from the sediment profiles, as well as directly dated bone harpoons. Three of the bones were also directly radiocarbon dated to the period approximately 3400-2800 BCE (Nielsen, 2020a, 2020b, 2020c; Nielsen & Persson, 2020).

All laboratory work prior to PCR was performed in a dedicated aDNA laboratory at the University of Oslo, following strict anti-contamination protocols (Gilbert et al., 2005; Llamas et al., 2017). Upon introduction to the aDNA lab, bones were brush-cleaned and UV-ed ten minutes on each side to reduce surface DNA contamination. The bones were then cut using an electric dentistry tool with an attached cutting disc in a sterile extraction hood, preserving morphometric landmarks. Cut fragments were crushed using a custom designed stainless-steel mortar as described in Gondek, Boessenkool, and Star (2018).

All samples were extracted using a standard extraction protocol adapted from Dabney et al. (2013) after a pre-digestion step (DD from Damgaard et al. 2015) or mild bleach treatment and pre-digestion (BleDD from Boessenkool et al., 2017) as described in Ferrari et al. (2021) (Table S1). In summary, powdered bone (2 × 200 mg per sample) was subjected to the DD or BleDD treatment and digested in 1 ml 0.5 M EDTA, 0.5 mg/ml proteinase K and 0.5% N-Laurylsarcosine for 18-24 h at 37 C. Combined digests were extracted with 9 × volumes of PB buffer (QIAGEN) and DNA was purified with MinElute columns on a QIAvac 24 Plus vacuum manifold system (QIAGEN).

Dual-indexed sequencing libraries were built as double stranded, blunt-ended libraries following Meyer and Kircher (2010) and Kircher et al. (2012) with modifications or as single stranded libraries following the Santa Cruz Reaction (SCR) protocol (Kapp et al.2021) (Table S1). Meyer and Kircher libraries were built from 20 μL of ligated DNA extract or extraction blanks and performed in half volumes reactions. The single stranded SCR libraries were built from 3-20 μL of ligated DNA extract (depending on the DNA concentration) or 20 μL extraction blanks using dilution tier 4. Indexing PCRs were performed with Taq Pfu Turbo Cx HotStart DNA polymerase (Agilent) with the following cycling conditions: 2 min activation at 95 C, 30 s denaturation at 95 C, 30 s annealing at 60 C, 1 min elongation at 72 C, and 10 min final extension at 72 C. Sample extracts were subject to 12 PCR cycles, while extraction blanks were subject to 30 PCR cycles to increase the chance of detecting contamination. Amplified libraries were cleaned using Agencourt AMPure XP PCR purification beads (Bronner et al., 2013) and examined on a Fragment Analyzer^TM^ (Advanced Analytical) using the High Sensitivity NGS Fragment Analysis Kit to determine suitability for sequencing. Libraries were sequenced on the Illumina HiSeq 4000 or NovaSeq 6000 (SP Flow Cell) platforms at the Norwegian Sequencing Centre with paired-end 150 bp reads and demultiplexed allowing zero mismatches in the index tag.

### Ancient specimens from the Mediterranean

92 individuals from archaeological excavations and zoological collections throughout the Mediterranean region dating from 100 to 1941 CE were obtained from Andrews et al. (in prep.) as BAM files (Table S2). These samples were prepared and extracted in the Ancient DNA Laboratory of the Department of Cultural Heritage (University of Bologna, Ravenna Campus, Italy), following strict criteria for aDNA analysis as per the Norwegian samples, and sequenced as single-stranded libraries (Kapp et al., 2021) at Macrogen facilities (Seoul, South Korea/Amsterdam, Netherlands) on a HiSeq X (100 bp paired-end) Illumina sequencing platform. Reads were processed using the Paleomix pipeline v.1.2.14 (Schubert et al., 2014) with settings described below (see “Bioinformatic processing of ancient and modern sequence data”), yielding an average of 28% endogenous DNA and 11-fold MT coverage (Table S8).

### Collection, extraction, and sequencing of modern samples

Modern tuna tissue samples of migratory, foraging adults (total-weight range: 151 - 313 kg) from Norway (NOR) (n = 38) were collected by the Norwegian Institute of Marine Research (IMR), from commercial catch off the coast of Møre og Romsdal, Western Norway. Two batches of modern Norwegian samples were obtained, the first from September 2018 and the second from September 2020 (Table S3). These batches were taken from two single purse seine catches and each of them are therefore assumed to belong to the same shoal. The 2018 batch of samples was freeze dried muscle tissue powdered at IMR facilities. The 2020 batch was collected as skin samples cut out between the spines of the dorsal fin and submerged immediately in RNAlater, shipped, and placed in a -20°C freezer within a week. The modern samples from Norway were all extracted in the modern DNA isolation laboratories at the University of Oslo, using the DNeasy Blood & Tissue kit (Qiagen) and following the manufacturer’s protocol.

Modern larvae or young-of-the-year (YoY) specimens (GOM: Gulf of Mexico, WMED: Western Mediterranean Balearic Islands, CMED: Central Mediterranean Sicily, EMED: Eastern Mediterranean Levantine Sea, n = 40, Table S4) were collected from each of the major Atlantic bluefin spawning sites (Figure 1) between 2013 and 2018. Juvenile albacore samples from the Bay of Biscay were caught by commercial vessels trolling in the Bay of Biscay between June and September of 2010 (Table S5). Larvae and tissue samples from each specimen were preserved in 96% ethanol and stored at -20 °C until further processing. Modern spawning site and albacore samples were extracted at the University of Bologna by a modified salt-based extraction protocol, as per Cruz et al. (2017), using SSTNE extraction buffer (Blanquer, 1990), and treated with RNase to remove residual RNA.

For all modern samples, DNA concentration was measured using a Qubit® dsDNA BR Assay Kit (Thermo Fisher Scientific, USA), where negative controls employed for each batch of samples extracted indicated undetectable levels of contamination. For the Norwegian samples, libraries were built using the TruSeq DNA Nano200 preparation kit (Illumina).

Modern spawning site extracts, along with albacore extracts, underwent single stranded library preparation following the SCR library protocol (Kapp et al., 2021). Sequencing and demultiplexing, allowing for zero mismatches, was performed at the Norwegian Sequencing Center on a combination of the HiSeq 4000 and NovaSeq 6000 (SP Flow Cell) Illumina sequencing platforms with paired-end 150 bp reads for all samples.

Raw sequence data of Pacific bluefin whole genome (Suda et al., 2019) were downloaded from DDBJ (accession no DRA008331) (Table S6) and used for interspecific population structure analyses.

### Bioinformatic processing of ancient and modern sequence data

Both modern and ancient reads were processed using the Paleomix pipeline v.1.2.14 (Schubert et al., 2014). Adapters were removed and forward and reverse reads were collapsed and trimmed with AdapterRemoval v.2.3.1 (Schubert et al., 2016), discarding collapsed reads shorter than 25 bp. All reads were aligned to a draft nuclear (NCBI BioProject: PRJNA408269) and MT reference genome (GenBank accession nr NC_014052.1) with BWA-mem v.0.7.17 (Li & Durbin, 2009) for mapping. All reads were filtered to a minimum Phred score quality of 25, so that only reads with higher mapping quality to the reference genome were considered endogenous and used for subsequent analyses. PCR duplicates were removed in Picard Tools v.2.18.27 and indel realignment (*GATKs IndelRealigner*) was performed to produce final BAM files. DNA post-mortem damage patterns were assessed in mapDamage v.2.0.9 (Ginolhac et al., 2011; Jónsson et al., 2013) after downsampling to 100,000 randomly selected reads.

MT BAMfiles were further processed in GATK v.4.1.4.0 following GATK best practices (McKenna et al., 2010). Individual genotypes were called (GATK v.4.1.4.0 *HaplotypeCaller -ploidy 1*) and then combined into a joint gvcf (GATK v.4.1.4.0 *CombineGVCFs*) before genotyping (GATK v.4.1.4.0 *GenotypeGVCFs*). Genotypes were hard-filtered in BCFtools v.1.9 (Li et al., 2009) (*-i ’FS<60.0 && SOR<4 && MQ>30.0 && QD > 2.0’ --SnpGap 10*) and VCFtools v.0.1.16 (Danecek et al., 2011) (*--minGQ 15 --minDP 2 --remove-indels*).

Filtered VCFs were indexed using Tabix v.0.2.6 (Li, 2011) and consensus sequences created as individual fasta files in BCFtools v.1.9 (bcftools consensus *-H 1*). Outgroup sequences were downloaded from GenBank (Clark et al., 2016) and curated using SeqKit v. 0.11.0 (*restart -i*) (Shen et al., 2016) so that all sequences started at position 1 in the D-loop, to correspond with the sample sequences. After renaming the fasta headers to their appropriate sample-IDs using BBMap v.38.50b (Bushnell, 2014) and combining the files to a multiple sequence alignment (MSA), the joint fasta files were aligned using MAFFT v.7.453 (Katoh et al., 2002) (*--auto*).

### Initial investigations and creation of datasets

Preliminary analyses were performed on a jointly called and filtered VCF and multiple-fasta, which included all samples. Missingness and depth was assessed for all samples using VCFtools v.0.1.16 (Danecek et al., 2011) and Principal Component analyses (PCA) (Adegenet (Jombart, 2008) in R v.4.1.2) and a Maximum Likelihood (ML) tree (IQTREE v. 1.6.12 (Nguyen et al., 2015), *-m MFP -bb 1000 -BIC*) was used to investigate clustering patterns and assess the presence of introgressed MT genomes. To look for identical haplotypes, we assessed the number of single nucleotide polymorphisms (SNPs) that differed between specimens using an in-house python3 script (https://github.com/laneatmore/nucleotide_differences), which uses MSAs as input to count the number of true SNP differences between all individuals (excluding those that were missing data) and generates distance matrices based on these differences.

Given that ancient bones may represent the same individual if they were obtained from the same archaeological context, we assessed the number of pairwise SNP differences between specimens at different filtering settings. To be conservative, ancient samples were considered identical if, in a pairwise comparison, they had no SNP differences at minDP2 *or* if they had only one SNP difference at minDP3 *and* came from the same archaeological layer. Using these criteria, we detected several genetically identical specimens of which the specimen with the highest endogenous DNA content was kept. Genetically identical modern samples were kept since distinct individuals were sampled. All samples with missingness above 50% (VCFtools v.0.1.16 *--missing-indv*, F_MISS) were also discarded. In total, 22 out of 208 samples (∼10%) were discarded from further analyses (Table S7 and S8).

Datasets were created for each sampling location depending on the presence of introgressed MT genomes (Table S13). First, a dataset excluding the introgressed individuals was created to allow for comparison of the effect of introgressed haplotypes on summary statistics. A haplotype was assumed to be introgressed if it clustered with albacore or Pacific bluefin in the PCA (Figure S3). This clustering was additionally supported by the ML and BEAST trees, which revealed the same individuals falling into highly supported monophyletic clades with the respective species (Figure S4). As the genotyping and filtering process in the GATK pipeline is affected by the haplotype variants present in the analyses, only samples within each dataset should be called and filtered together to accurately present the variation (GATK, 2016). These separate datasets were therefore separately genotyped, filtered, and aligned (with settings described in the section above) to create respective multiple-fasta files for subsequent analyses. An overview of these individual datasets can be found in Table S14.

### Population genomic analyses

Genetic population structure was investigated using Principal Component analyses (PCA) as implemented in the R-package Adegenet (Jombart, 2008). A map of missing loci and alleles diverging from the reference genome, was created to assess missing genotypes in both ancient and modern samples and better visualize introgressed specimens. All plots were created with R 4.3 in RStudio (Rstudio Team 2021), using various packages for data loading, analyses, and visualization (see supplementary Section 1).

Genetic diversity was investigated using a range of standard population genetic measurements. The number of haplotypes (Nh) and haplotype diversity (hD) (Nei 1987) was calculated in R-package pegas (Paradis, 2010) and independently assessed in Fitchi (Matschiner, 2016). The number of segregating sites (S), nucleotide diversity (π) (Nei 1987), Tajima’s D (TD) (Tajima, 1989), and Fu and Li’s F statistic (F) (Fu & Li, 1993) were calculated in DnaSP v.6 (Rozas et al., 2017).S, π, TD values were confirmed in pegas. To account for differences in sample sizes across sites when calculating π and TD, an additional analysis using 1000 bootstrap replicates and subsampling five individuals per round without replacement, was performed in pegas on datasets where the total sample size was over five.

Phylogenetic relationships were investigated using both ML and Bayesian approaches. ML trees with 100 nonparametric bootstrap replicates were created in IQTREE v. 1.6.12 (Nguyen et al., 2015). ModelFinder Plus (MFP) (Kalyaanamoorthy et al., 2017) was used to search all available models, and best-fit models were selected according to the Bayesian Information Criterion (BIC) (Schwarz, 1978). Bayesian trees were created in BEAST 2 v.2.6.4 (R. Bouckaert et al., 2014), using the Yule model prior under a strict clock with mutation rate 3.6×10^-8^ substitutions per site per year as per Donaldson and Wilson (1999), running MCMC over 800,000,000 generations and sampling once every 1000 generations. bModelTest (R. R. Bouckaert & Drummond, 2017) was used to assess available site models, and the resulting logfile was inspected in Tracer v1.7.2 (Rambaut et al., 2018). Trees were downsampled in LogCombiner (implemented in BEAST 2 v.2.6.4), resampling every 10,000 trees. TreeAnnotator (implemented in BEAST 2 v.2.6.4) was used to remove the first 10% of the trees (burnin) and create a target maximum clade credibility tree. Nodes with less than 50% posterior support were excluded from the summary analysis in TreeAnnotator so that only nodes present in the majority of the trees were annotated. The final trees in all phylogenetic analyses were visualized and curated in FigTree v.1.4.4 (Rambaut, 2018).

Evolutionary relationships were visualized using haplotype networks created in Fitchi (*--haploid -p*) (Matschiner, 2016) using the ML trees generated in IQTREE (described above) as input with bootstrap values removed using the R-package ape (Paradis et al., 2004). For the dataset including introgressed individuals and outgroup species, a minimum edge length of seven substitutions was defined (*-e 7*) so that haplotypes separated by seven or fewer substitutions were collapsed into one node. For the dataset only containing Atlantic bluefin haplotypes, each node was defined as a unique haplotype (*-e 1*).

Genetic distance between sample locations was assessed using measures of absolute (dxy) and relative (ΦST) divergence, calculated using DnaSP v.6 (“*DNA divergence between populations”, all sites*) (Rozas et al., 2017) and Arlequin v.3.5 (Excoffier & Lischer, 2010) respectively. In Arlequin, pairwise ΦST was calculated via a distance matrix computed by Arlequin based on Tamura & Nei (1993) and assuming no rate heterogeneity, as suggested by bModelTest (R. R. Bouckaert & Drummond, 2017) (implemented in BEAST 2 v.2.6.4 (R. Bouckaert et al., 2014)). To test the significance of ΦST, p-values were generated in Arlequin using 1000 permutations.

## Results

### DNA yield and library success

A total of 1.7 billion sequencing reads were obtained for the 38 ancient samples from Norway. These specimens had remarkable DNA preservation with 100% library success and yielding, on average, 24% endogenous DNA and 20-fold MT coverage (Table S7). The reads showed postmortem degradation patterns expected for authentic ancient DNA (Figure S2). A total of 3.1 billion sequencing reads were obtained for the 84 modern specimens, resulting in 711-fold MT coverage on average for the 78 Atlantic bluefin specimens (Table S9 and S10) and 221-fold MT coverage on average for the six albacore samples (Table S11). The Pacific bluefin raw sequence data from Suda et al. (2019) yielded 3322-fold MT coverage (Table S12). After stringent filtering, 186 out of 208 specimens (∼90%) were kept for further analyses (Table S7 and S8).

### Detecting introgressed MT mitogenomes

Out of 186 samples analyzed, seven ancient and four modern individuals had MT haplotypes that clustered closely with albacore or Pacific bluefin in both the PCA, haplotype network and phylogenies. The PCA reveals three distinct clusters (Figure 2) with PC1 separating an Atlantic bluefin cluster from a Pacific bluefin and albacore cluster and PC2 separating the latter two speciese. Within the Pacific bluefin cluster, we observe two modern (both NOR) and four ancient (two Norway 3000BCE, one Istanbul 800-1200CE and one Sardinia 1500-1700CE) Atlantic bluefin specimens. Within the albacore cluster, we observe two modern (one NOR and one WMED) and three ancient (one Istanbul-800-1200CE and two Sicily-900-1200CE) Atlantic bluefin specimens. The PCA-clusters are reiterated in the haplotype network (Figure 2). ML and Bayesian phylogenetic analyses provided full statistical support (bootstrap=100, posterior probability=1) for the three species as monophyletic groups with the same six Pacific-like and five albacore-like haplotypes again clustering with their respective species (Figure 2, see also Figure S6). We conclude that these Pacific-like and albacore-like haplotypes are introgressed MT genomes into Atlantic bluefin tuna.

**Figure 2:**
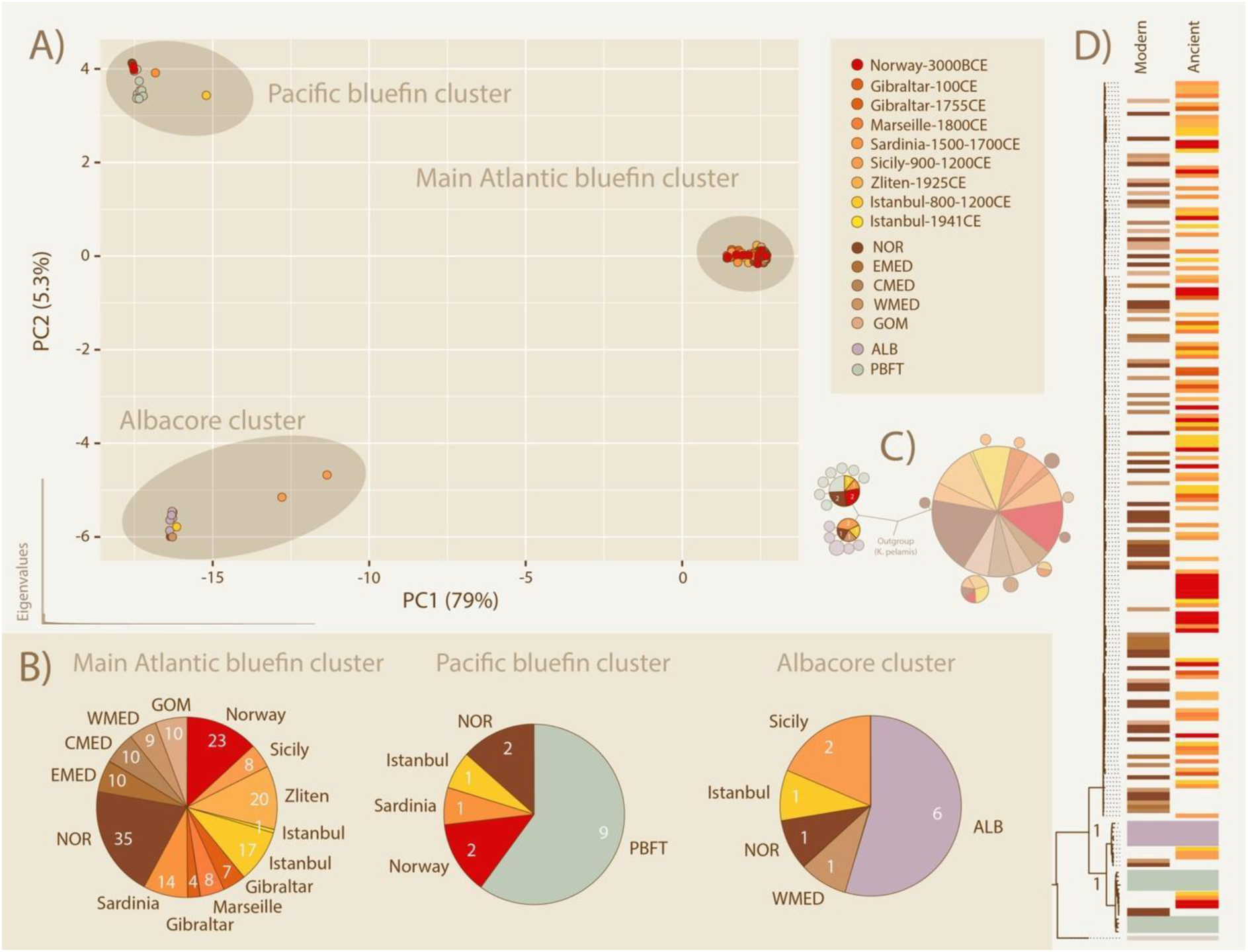
Species clusters and introgressed MT haplotypes within Atlantic bluefin specimens, revealed by PCA, haplotype network and phylogenetic analyses A) Three species specific clusters detected in ancient and modern Atlantic tuna specimens. The PCA shows three species specific clusters and PCA eigenvalues are shown in the bottom left corner. Modern modern Pacific tuna (blue) and albacore (grey) specimens are included as controls. B) Relative abundance of haplotypes per location within each PCA-cluster is visualized as pie-charts, with the number of samples from each location indicated on the slices. C) Haplotype network showing three species specific haplotypes. Haplotypes separated by seven or fewer substitutions were collapsed into single nodes. D) Interspecific phylogeny of specimens with posterior probability support for the species clades (see also Figure S6). Colors are representative of the spatiotemporal cohorts listed in the legend of panel A).

### Spatiotemporal population structure

We find no significant genomic differentiation between any of the temporal cohorts. We also observe no spatial differences in the level of genetic variation between any of the sampling locations for Atlantic bluefin. Atlantic bluefin individuals from both management stocks and across the Eastern stock range and spawning areas are scattered across the intraspecific PCA (Figure S8) and haplotype network (Figure S9). The sampling locations are also distributed along the entire phylogeny within the Atlantic bluefin group (Figure 2, see also Figure S6 and S10). The intraspecific Atlantic bluefin haplotype network reveals a star-like pattern with more recent haplotypes deriving from an ancestral, central haplotype (Figure S9).

### Genetic divergence and diversity influenced by introgression

Measures of pairwise genetic distance between Atlantic bluefin sampling locations show low levels of absolute (dxy) and relative (Φst) divergence, either excluding (Figure 3a) or including introgressed MT haplotypes (Figure 3b). Genetic differentiation increases when introgressed samples are included. This increase is driven by the inclusion of genetically more divergent haplotypes, which are not present at each location or temporal cohort. In all cases, levels of Φst remained low and non-significant (Figure S7) across all populations. Including individuals with introgressed MT haplotypes increased values of nucleotide diversity π and S (Table S15). The number of haplotypes (hD) is not impacted; most sample locations only contained unique specimens (N=Nh) therefore leading to a hD of 1, meaning 100% probability of obtaining unique samples during random sampling. Tajima’s D (TD) was also not affected by the inclusion of introgressed individuals and was significantly negative for most locations and temporal cohorts and when analyzing all specimens jointly (Table S15).

**Figure 3:**
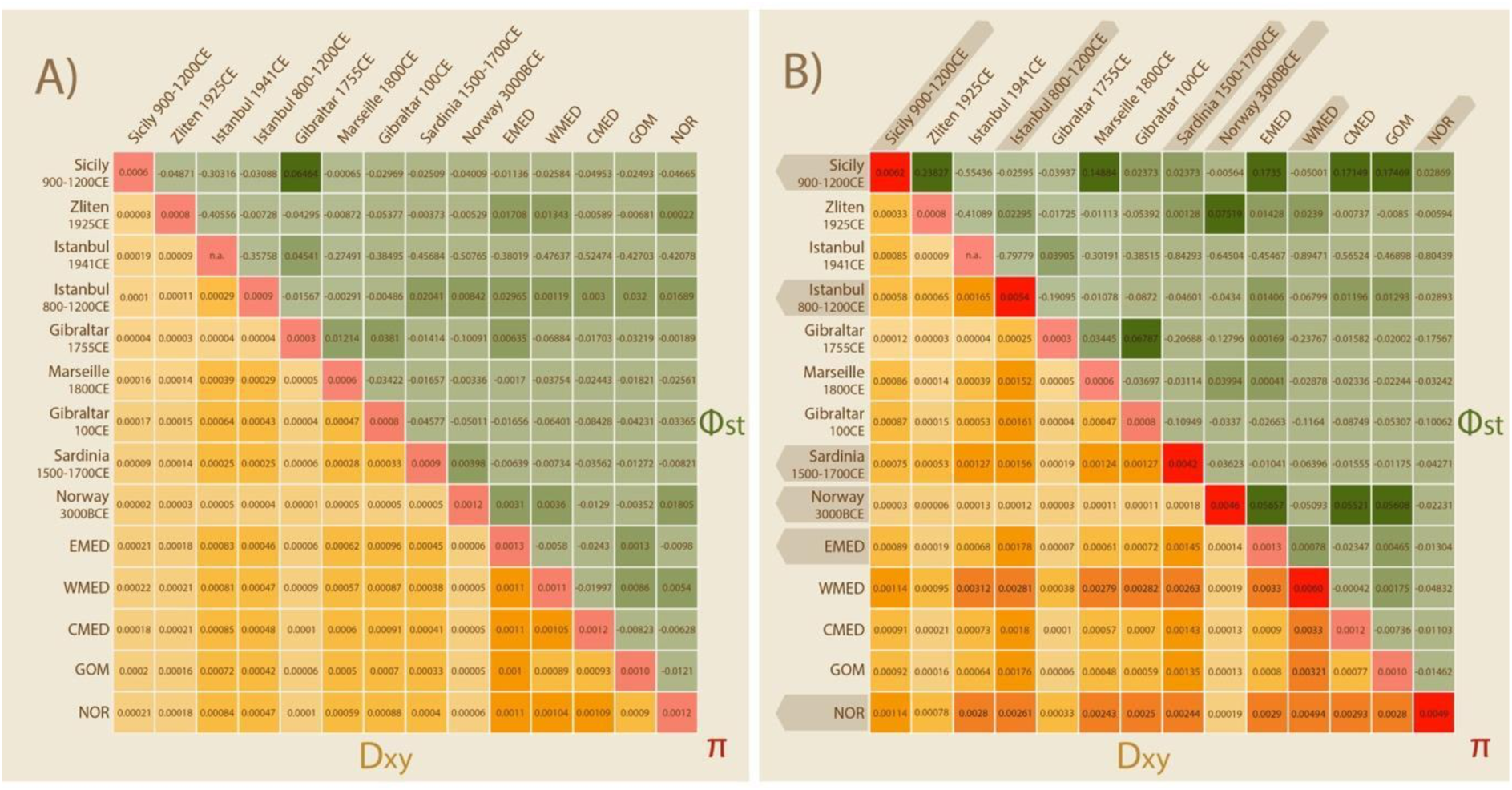
No significant spatiotemporal population structure in Atlantic bluefin tuna based on mitogenomic data of 186 specimens. Pairwise population divergence is presented as a heatmap showing absolute (dxy) and relative (ΦST) divergence between populations when A) excluding and B) including the introgressed mitogenomes. Divergence is inflated when including introgressed mitogenomes. Locations containing introgressed individuals are highlighted with darker shading in panel B). The nucleotide diversity within each population is shown on the diagonal. P-values for ΦST can be found in supplementary (Figure S7).

### MT haplotype introgression over time

We observe MT haplotype introgression into Atlantic bluefin throughout a 5000-year chronology (Figure 4). The earliest observation is the presence of two introgressed MT haplotypes from Pacific bluefin in the Neolithic (ca. 3000 BCE) in Norway. Pacific bluefin tuna MT haplotypes are further found in early medieval Istanbul (800-1200CE), late-medieval Sardinia (1500-1700 CE) and modern Norway. Albacore MT haplotypes are found in early medieval Istanbul (800-1200CE) and Sicily (900-1200CE), modern Western Mediterranean and modern Norway.

**Figure 4:**
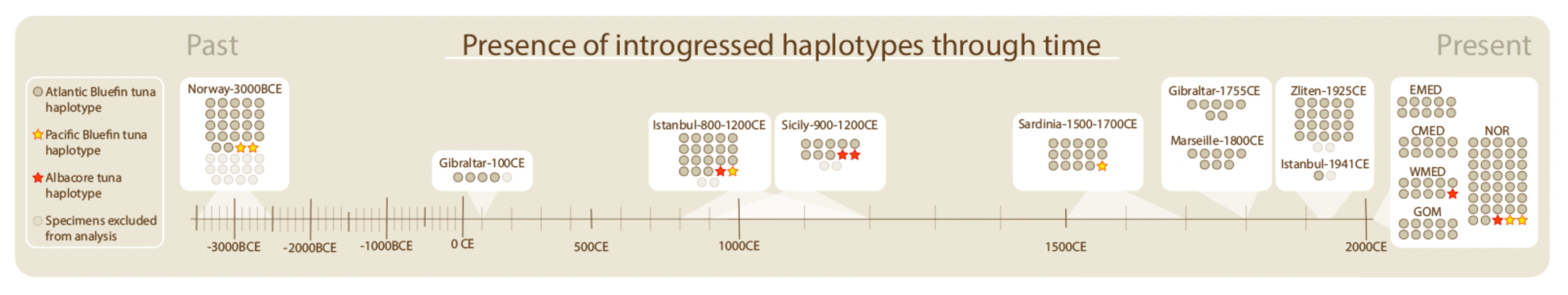
Introgressed Pacific bluefin MT haplotypes (yellow stars) or albacore MT haplotypes (red stars) are observed along an entire 5000-year-old chronology of ancient Atlantic bluefin tuna specimens. Individual tuna specimens (n=208) (circle or star) are grouped according to their age, determined by archaeological context. Specimens were either modern (n=78), or ancient (n=130) and dated by archaeological context. Uncertainty in the age range of ancient specimens is depicted beneath their respective sample sets (light shading).

## Discussion

We here present a 5000-year chronology of MT introgression in the Atlantic bluefin tuna. The presence of introgressed MT haplotypes in the Neolithic (ca. 3000 BCE), turn of the millennium (800-1200 CE), and in present day populations reveals that introgression resulted from long-term natural phenomena rather than being a consequence of recent anthropogenic impacts. Moreover, our results show that the frequency of introgression in the Atlantic bluefin has remained stable over millennia despite shifts in abundance and distribution of Atlantic bluefin populations.

### Evidence of mitochondrial introgression through time

We obtain similar rates of MT introgression as reported by previous studies in our modern samples, with rates of introgression for Pacific bluefin and albacore at 2.6% (2/78 individuals) each. Including our ancient samples in the calculation, the rate of introgression remains astonishingly stable with 3.2% Pacific bluefin haplotypes (6/186) and 2.7% albacore-like haplotypes (5/186) across all samples. In total, six Pacific-like and five albacore-like haplotypes consistently cluster with their respective species through all interspecific analyses. We do not observe albacore-like haplotypes from the Neolithic period, but because of the low frequency of introgression compared to the sample size we speculate this is likely due to sampling stochasticity rather than lack of natural hybridization at the time. Still, it cannot be excluded that Neolithic climate conditions drove albacore populations away from areas inhabited by Atlantic bluefin.

The location of the hybridization events with introgressive hybridization from both albacore and Pacific bluefin remain to be investigated. While the albacore has overlapping ranges and spawning areas with both bluefin species, the Pacific and Atlantic bluefins are geographically separated with no documented migration. The potential migration of Pacific bluefin into the Atlantic Ocean has been hypothesized to occur via the Indian Ocean and following the Agulhas current around the tip of Africa (Bremer et al. 2005). Whether this represents a contemporary migration route or a historical process where past, stronger currents might have facilitated admixture is unclear (Bremer et al. 2005). The similar rate of introgression from both albacore and Pacific bluefin and the lack of introgressed MT haplotypes in the albacore makes the range-overlapping albacore an unlikely carrier of MT haplotypes between the bluefins. As Pacific bluefin also contains Atlantic bluefin-like MT haplotypes, another hypothesis for their presence is incomplete lineage sorting (ILS). The Thunnus genus is thought to have diverged rapidly within the last 6-10 million years, with a more recent speciation of the Pacific and Atlantic bluefins only around 400,000 years ago (Santini et al. 2013; Ciezarek et al. 2018; Díaz-Arce et al. 2016). While introgressive hybridization is the likely origin of albacore-like haplotypes in both bluefin species (Ciezarek et al. 2018), observed gene-tree versus species-tree discordance that did not deviate from expectations under ILS. These results indicate that the observed patterns of Pacific-like MT haplotypes in the Atlantic bluefin population, and vice versa, may be a result of ILS rather than introgressive hybridization. Larger genomic databases are required to furhter delineate between these two hypotheses.

While our historical investigation focuses on the eastern Atlantic with samples from the Mediterranean and Norway, introgressed MT genomes from albacore have also been found in the Gulf of Mexico (1% frequency) and the Slope Sea (6% frequency). Because introgressed MT genomes in Atlantic bluefin were first observed in the eastern Atlantic, their presence in the western Atlantic has been hypothesized to be introduced via gene flow from the Mediterranean (Diaz-Arce et al., 2024). An increase in gene flow from the Mediterranean into the Gulf of Mexico and Slope Sea will likely erode genetic differences between the two management stocks (Diaz-Arce et al., 2024). Direct observations of intogression events within the Western Atlantic bluefin stock have not been made, although albacore is known to spawn across tropical waters including the South-West Sargasso Sea as well as the Mediterranean (ICCAT, 2016, 2020; NOAA, 2023). Future studies should investigate long-term MT introgression in the Gulf of Mexico to disentangle the recently suggested changes in demographic patterns (Diaz-Arce et al., 2024) and locate the origins of the intogression events.

### Introgressed haplotypes impact estimates of genomic differentiation

We observe that the presence of introgressed haplotypes increases measures of genetic diversity (S and π), which is driven by the highly divergent alleles of such introgressed MT genomes (Figure S5). The inclusion of these diverging haplotypes also influences the measures of genetic divergence (dxy and ΦST) between locations and temporal cohorts, consistently increasing the absolute genetic diversity (dxy) and altering the pattern of relative genetic divergence (ΦST). Tajima’s D (TD) was not consistently affected by the inclusion of introgressed individuals, although in some cases TD changed value and lost or attained significance when introgressed haplotypes were included (Table S15). Considering these results, it is important to be aware of highly diverging haplotypes when extrapolating population genetic statistics from subsamples of natural populations containing highly diverging haplotypes. The low frequency of natural hybridization in Atlantic bluefin causes stochasticity at low sample sizes, and we find that the inclusion of introgressed individuals inflates population genomic statistics that are commonly used for management and population viability assessments (Hohenlohe et al., 2021).

### Spatiotemporal population structure

We find no significant divergence and no pattern of genomic MT differentiation between any of the spatial or temporal cohorts. Ancient and modern samples largely intermixed in all analyses, suggesting mitogenomic stability and temporal continuity through time. Similar observations in other species, such as Atlantic cod (*Gadus morhua*) (Martínez-García et al., 2021) and New Zealand snapper (*Chrysophrys auratus*) (Oosting et al., 2023) emphasize the low power of the MT genome to observe spatiotemporal differentiation in wide ranging fish species. The regular presence of identical haplotypes across sampling locations and temporal cohorts emphasizes the lack of mitogenomic variation and informative markers for population structure in this species. Although we cautiously removed identical samples from the same archaeological excavations, the presence of identical samples across cohorts that were processed in different laboratories shows this as a naturally occurring phenomenon.

Population genetic statistics confirmed low mitogenomic variation with no significant divergence between any of the spatiotemporal cohorts and no temporal loss of genetic diversity despite heavy exploitation (Figure 3A, Table S15). Across datasets, TD was negative and often significant, suggesting an excess of rare alleles in the datasets. This is indicative of either positive selection or recent population expansion (Fijarczyk & Babik, 2015; Delph & Kelly, 2014). Population expansion is further corroborated in the intraspecific haplotype network, where newer haplotypes are derived from a shared central haplotype forming a star-like pattern (Figure S9). These results highlight robust preservation of the MT genome despite centuries of human exploitation.

## Conclusion

Atlantic bluefin tuna has experienced significant changes in distribution linked to sea surface temperature oscillations during the past centuries (Faillettaz et al., 2019; Muhling et al., 2011; Ravier & Fromentin, 2004), alongside intense exploitation (Andrews et al., 2022; Block, 2019), biomass depletion, range contraction, trophic niche loss (Andrews et al., 2023; Di Natale, 2015; Tangen, 2009), followed by recovery, increased biomass, and range expansion of the Eastern Atlantic bluefin stock during the last decade (Nøttestad et al. 2020; ICCAT 2023). Despite such extensive spatial shifts in distribution over time, we show that MT introgression is a long-term natural phenomenon in Atlantic bluefin. The stable frequency over time suggests that this phenomenon is robust against historical climate fluctuations and more recent anthropogenic impacts. By providing a baseline observation, our study highlight the utility of ancient DNA to obtain temporal insights in the long-term persistence of such phenomena, which is essential for disentangling the drivers of introgression and any fitness consequences of MT intogression.

## Supporting information

ENA-sample-bamfiles

## Acknowledgements

This work was supported by the European Union’s Horizon 2020 Research and Innovation Programme under the Marie Skłodowska-Curie grant agreement No. 813383 (SeaChanges) and the 4-OCEANS Synergy grant agreement no. 951649. The European Research Agency is not responsible for any use that may be made of the information this work contains. Special thanks goes to the Norwegian Sequencing Centre (University of Oslo; https://www.sequencing.uio.no) for the sequencing of genomic libraries. Computation was performed using the resources and assistance from SIGMA2. The collection of recent samples in Norway was financed by the Institute of Marine Research. The collection of larva samples in the Gulf of Mexico was funded by the NOAA RESTORE Science Program, BLOOFINZ-GoM #NOAA-NOS-NCCOS-2017-2004875; Cooperative Institute for Marine and Atmospheric Studies #NA20OAR4320472. This work is a contribution to the https://tunaarchaeology.org/ project.

We are grateful to Ørjan Sørensen at the Institute of Marine Research (IMR) for his contributions to the collection and shipping of the modern Atlantic bluefin samples from Norway and providing sample metadata. We thank Verónica Gómez-Fernández (Instituto Nacional de Investigaciones Científicas y Ecológicas, Spain), Gabriele Carenti (CEPAM, CNRS, Université Côte d’Azur, France) and Darío Bernal-Casasola (Department of History, Geography and Philosophy, Faculty of Philosophy and Letters, University of Cádiz, Spain) for sample collection. We thank Leif Nøttestad (IMR) and Mark Ravinet (CEES, University of Oslo) for comments and stimulating scientific discussions.

## Author contributions following CRediT (https://credit.niso.org/)

Emma Falkeid Eriksen: Conceptualization, Data curation, Formal Analysis, Investigation, Methodology, Visualization, Writing – original draft, Writing – review & editing

Adam Jon Andrews: Data curation, Investigation, Methodology, Writing – review & editing Svein Vatsvåg Nielsen: Resources, Writing – review & editing

Per Persson: Writing – review & editing

Estrella Malca: Resources, Writing – review & editing Vedat Onar: Resources

Veronica Aniceti: Resources Gäel Piquès: Resources

Federica Piattoni: Data curation, methodology, Writing – review & editing Francesco Fontani: Investigation

Martin Wiech: Resources, Writing-review & editing Keno Ferter: Resources, Writing-review & editing Oliver Kersten: Investigation, Writing-review & editing Giada Ferrari: Investigation, Writing-review & editing

Alessia Cariani: Resources, Funding acquisition, Writing-review & editing Fausto Tinti: Resources, Funding acquisition, Writing-review & editing Elisabetta Cilli: Resources, Funding acquisition

Lane M. Atmore: Software, Supervision, Writing – review & editing

Bastiaan Star: Conceptualization, Funding acquisition, Project administration, Software, Supervision, Writing – original draft, Writing – review & editing

## Data archiving

All mitochondrial BAM files used in this study has been uploaded in the European Nucleotide Archive (ENA) and can be accessed at https://www.ebi.ac.uk/ena/browser/view/PRJEB74135. See supplementary csv containing filenames, project accession number and ENA identifier for all samples.

## Supplementary

### Supplementary Section 1

R-packages used in population genomic analyses

- vcfR (data loading) (Knaus & Grünwald, 2017)
- adegenet (ordination analysis) (Jombart, 2008)
- ape (phylogenetic analyses) (Paradis et al., 2004)
- pegas (population genomic statistics) (Paradis, 2010)
- ggplot in tidyverse (visualization) (Wickham et al., 2019)
- gridExtra (visualization) (Auguie and Antonov 2017)
- lemon (visualization) (Edwards 2017)

### Section 2: Supplementary Tables

**Table S1:** Sample locations, laboratory protocols and dsDNA concentration for the ancient Atlantic bluefin samples from Norway. All samples were taken from bone tissue (vertebrae) and all samples were dated by context to be from 3000 BCE. More information about the sites can be found in the archaeological reports (Nielsen, 2020a, 2020b, 2020c).

| Sample-ID | Coordinates | Extraction protocol | Qubit concentration | Library protocol |
| --- | --- | --- | --- | --- |
| aTUNn02-Norway-3000BCE | N 58.28 E 8.50 | DD | 1.24 | M&K DS |
| aTUNn03-Norway-3000BCE | N 58.28 E 8.50 | DD | 0.48 | M&K DS |
| aTUNn04-Norway-3000BCE | N 58.28 E 8.50 | DD | 1.27 | M&K DS |
| aTUNn05-Norway-3000BCE | N 58.28 E 8.50 | DD | 1.02 | M&K DS |
| aTUNn06-Norway-3000BCE | N 58.28 E 8.50 | DD | 2.7 | M&K DS |
| aTUNn07-Norway-3000BCE | N 58.28 E 8.50 | bleDD | n.a. | M&K DS |
| aTUNn08-Norway-3000BCE | N 58.28 E 8.50 | bleDD | n.a. | M&K DS |
| aTUNn09-Norway-3000BCE | N 58.28 E 8.50 | bleDD | n.a. | M&K DS |
| aTUNn10-Norway-3000BCE | N 58.28 E 8.50 | bleDD | n.a. | M&K DS |
| aTUNn11-Norway-3000BCE | N 58.28 E 8.50 | bleDD | n.a. | M&K DS |
| aTUNn12-Norway-3000BCE | N 58.28 E 8.50 | bleDD | n.a. | M&K DS |
| aTUNn13-Norway-3000BCE | N 58.28 E 8.50 | bleDD | n.a. | M&K DS |
| aTUNn14-Norway-3000BCE | N 58.28 E 8.50 | bleDD | n.a. | M&K DS |
| aTUNn15-Norway-3000BCE | N 58.28 E 8.50 | bleDD | n.a. | M&K DS |
| aTUNn16-Norway-3000BCE | N 58.28 E 8.50 | bleDD | n.a. | M&K DS |
| aTUNn17-Norway-3000BCE | N 58.28 E 8.50 | bleDD | 1.28 | SC SS |
| aTUNn18-Norway-3000BCE | N 58.28 E 8.50 | bleDD | 13.3 | SC SS |
| aTUNn19-Norway-3000BCE | N 58.28 E 8.50 | bleDD | 5.6 | SC SS |
| aTUNn20-Norway-3000BCE | N 58.28 E 8.50 | bleDD | 1.6 | SC SS |
| aTUNn21-Norway-3000BCE | N 58.28 E 8.50 | bleDD | 2.8 | SC SS |
| aTUNn22-Norway-3000BCE | N 58.28 E 8.50 | bleDD | 5.56 | SC SS |
| aTUNn23-Norway-3000BCE | N 58.28 E 8.50 | bleDD | 19.6 | SC SS |
| aTUNn24-Norway-3000BCE | N 58.28 E 8.50 | bleDD | 3.54 | SC SS |
| aTUNn25-Norway-3000BCE | N 58.28 E 8.50 | bleDD | 11 | SC SS |
| aTUNn26-Norway-3000BCE | N 58.28 E 8.50 | bleDD | 6.94 | SC SS |
| aTUNn27-Norway-3000BCE | N 58.28 E 8.50 | bleDD | 2.8 | SC SS |
| aTUNn28-Norway-3000BCE | N 58.28 E 8.50 | bleDD | 10.9 | SC SS |
| aTUNn29-Norway-3000BCE | N 58.28 E 8.50 | bleDD | 89.6 | SC SS |
| aTUNn30-Norway-3000BCE | N 58.28 E 8.50 | bleDD | 1.87 | SC SS |
| aTUNn31-Norway-3000BCE | N 58.28 E 8.50 | bleDD | 19.9 | SC SS |
| aTUNn32-Norway-3000BCE | N 58.28 E 8.50 | bleDD | 26 | SC SS |
| aTUNn33-Norway-3000BCE | N 58.28 E 8.50 | bleDD | 1.9 | SC SS |
| aTUNn34-Norway-3000BCE | N 58.28 E 8.50 | bleDD | 36 | SC SS |
| aTUNn35-Norway-3000BCE | N 58.28 E 8.50 | bleDD | 78.8 | SC SS |
| aTUNn36-Norway-3000BCE | N 58.28 E 8.50 | bleDD | 25.2 | SC SS |
| aTUNn37-Norway-3000BCE | N 58.28 E 8.50 | bleDD | 3.18 | SC SS |
| aTUNn38-Norway-3000BCE | N 58.28 E 8.50 | bleDD | 3.06 | SC SS |
| aTUNn39-Norway-3000BCE | N 58.28 E 8.50 | bleDD | 1.4 | SC SS |

DD: Double digestion. Extraction protocol adapted from Damgaard et al. 2015.
bleDD: Bleach and double digestion. Extraction protocol adapted from Boessenkool et al. 2017.
M&K DS: Double stranded libraries. Library protocol adapted from Meyer and Kircher 2010 with modifications from Schroeder et al. 2015.
SC SS: Santa Cruz Reaction single-stranded library protocol from Kapp, Green, and Shapiro 2021.

**Table S2:** Sample locations, date sampled and for the ancient Atlantic bluefin samples from the Mediterranean. All samples were taken from bone tissue (vertebrae). Samples were dated by context at the archaeological sites. More information about the sites can be found in Andrews et al. (2023b).

| Sample-ID | Coordinates | Location | Date of origin |
| --- | --- | --- | --- |
| aTUNm01-Sicily-900-1200CE | N 38.11 E 13.37 | Palermo | 900-1000 CE |
| aTUNm02-Sicily-900-1200CE | N 38.11 E 13.37 | Palermo | 900-1000 CE |
| aTUNm03-Sicily-900-1200CE | N 38.11 E 13.37 | Palermo | 900-1000 CE |
| aTUNm04-Sicily-900-1200CE | N 38.11 E 13.37 | Palermo | 900-1000 CE |
| aTUNm05-Sicily-900-1200CE | N 38.11 E 13.37 | Palermo | 900-1000 CE |
| aTUNm06-Sicily-900-1200CE | N 38.11 E 13.37 | Palermo | 900-1000 CE |
| aTUNm07-Sicily-900-1200CE | N 37.65 E 12.59 | Mazara del Vallo | 1200 CE |
| aTUNm08-Sicily-900-1200CE | N 37.65 E 12.59 | Mazara del Vallo | 1200 CE |
| aTUNm09-Sicily-900-1200CE | N 37.65 E 12.59 | Mazara del Vallo | 1200 CE |
| aTUNm10-Sicily-900-1200CE | N 37.65 E 12.59 | Mazara del Vallo | 1200 CE |
| aTUNm11-Sicily-900-1200CE | N 38.11 E 13.36 | Palermo | 900-1000 CE |
| aTUNm12-Sicily-900-1200CE | N 38.11 E 13.36 | Palermo | 900-1000 CE |
| aTUNm13-Zliten-1925CE | N 33.25 E 14.66 | Zliten | 1925 CE |
| aTUNm14-Zliten-1925CE | N 33.25 E 14.66 | Zliten | 1925 CE |
| aTUNm15-Zliten-1925CE | N 33.25 E 14.66 | Zliten | 1925 CE |
| aTUNm16-Zliten-1925CE | N 33.25 E 14.66 | Zliten | 1925 CE |
| aTUNm17-Zliten-1925CE | N 33.25 E 14.66 | Zliten | 1925 CE |
| aTUNm18-Zliten-1925CE | N 33.25 E 14.66 | Zliten | 1925 CE |
| aTUNm19-Zliten-1925CE | N 33.25 E 14.66 | Zliten | 1925 CE |
| aTUNm20-Zliten-1925CE | N 33.25 E 14.66 | Zliten | 1925 CE |
| aTUNm21-Zliten-1925CE | N 33.25 E 14.66 | Zliten | 1925 CE |
| aTUNm22-Zliten-1925CE | N 33.25 E 14.66 | Zliten | 1925 CE |
| aTUNm23-Zliten-1925CE | N 33.25 E 14.66 | Zliten | 1925 CE |
| aTUNm24-Zliten-1925CE | N 33.25 E 14.66 | Zliten | 1925 CE |
| aTUNm25-Zliten-1925CE | N 33.25 E 14.66 | Zliten | 1925 CE |
| aTUNm26-Zliten-1925CE | N 33.25 E 14.66 | Zliten | 1925 CE |
| aTUNm27-Zliten-1925CE | N 33.25 E 14.66 | Zliten | 1925 CE |
| aTUNm28-Zliten-1925CE | N 33.25 E 14.66 | Zliten | 1925 CE |
| aTUNm29-Zliten-1925CE | N 33.25 E 14.66 | Zliten | 1925 CE |
| aTUNm30-Zliten-1925CE | N 33.25 E 14.66 | Zliten | 1925 CE |
| aTUNm31-Zliten-1925CE | N 33.25 E 14.66 | Zliten | 1925 CE |
| aTUNm32-Zliten-1925CE | N 33.25 E 14.66 | Zliten | 1925 CE |
| aTUNm33-Zliten-1925CE | N 33.25 E 14.66 | Zliten | 1925 CE |
| aTUNm34-Zliten-1925CE | N 33.25 E 14.66 | Zliten | 1925 CE |
| aTUNm35-Istanbul-1941CE | N 41.01 E 28.95 | Istanbul | 1941 CE |
| aTUNm36-Istanbul-1941CE | N 41.01 E 28.95 | Istanbul | 1941 CE |
| aTUNm37-Istanbul-800-1200CE | N 41.01 E 28.95 | Istanbul | 800-1200 CE |
| aTUNm38-Istanbul-800-1200CE | N 41.01 E 28.95 | Istanbul | 800-1200 CE |
| aTUNm39-Istanbul-800-1200CE | N 41.01 E 28.95 | Istanbul | 800-1200 CE |
| aTUNm40-Istanbul-800-1200CE | N 41.01 E 28.95 | Istanbul | 800-1200 CE |
| aTUNm41-Istanbul-800-1200CE | N 41.01 E 28.95 | Istanbul | 800-1200 CE |
| aTUNm42-Istanbul-800-1200CE | N 41.01 E 28.95 | Istanbul | 800-1200 CE |
| aTUNm43-Istanbul-800-1200CE | N 41.01 E 28.95 | Istanbul | 800-1200 CE |
| aTUNm44-Istanbul-800-1200CE | N 41.01 E 28.95 | Istanbul | 800-1200 CE |
| aTUNm45-Istanbul-800-1200CE | N 41.01 E 28.95 | Istanbul | 800-1200 CE |
| aTUNm46-Istanbul-800-1200CE | N 41.01 E 28.95 | Istanbul | 800-1200 CE |
| aTUNm47-Istanbul-800-1200CE | N 41.01 E 28.95 | Istanbul | 800-1200 CE |
| aTUNm48-Istanbul-800-1200CE | N 41.01 E 28.95 | Istanbul | 800-1200 CE |
| aTUNm49-Istanbul-800-1200CE | N 41.01 E 28.95 | Istanbul | 800-1200 CE |
| aTUNm50-Istanbul-800-1200CE | N 41.01 E 28.95 | Istanbul | 800-1200 CE |
| aTUNm51-Istanbul-800-1200CE | N 41.01 E 28.95 | Istanbul | 800-1200 CE |
| aTUNm52-Istanbul-800-1200CE | N 41.01 E 28.95 | Istanbul | 800-1200 CE |
| aTUNm53-Istanbul-800-1200CE | N 41.01 E 28.95 | Istanbul | 800-1200 CE |
| aTUNm54-Istanbul-800-1200CE | N 41.01 E 28.95 | Istanbul | 800-1200 CE |
| aTUNm55-Istanbul-800-1200CE | N 41.01 E 28.95 | Istanbul | 800-1200 CE |
| aTUNm56-Istanbul-800-1200CE | N 41.01 E 28.95 | Istanbul | 800-1200 CE |
| aTUNm57-Istanbul-800-1200CE | N 41.01 E 28.95 | Istanbul | 800-1200 CE |
| aTUNm58-Gibraltar-1755CE | N 36.28 W 6.09 | Conil | 1755 CE |
| aTUNm59-Gibraltar-1755CE | N 36.28 W 6.09 | Conil | 1755 CE |
| aTUNm60-Gibraltar-1755CE | N 36.28 W 6.09 | Conil | 1755 CE |
| aTUNm61-Gibraltar-1755CE | N 36.28 W 6.09 | Conil | 1755 CE |
| aTUNm62-Gibraltar-1755CE | N 36.28 W 6.09 | Conil | 1755 CE |
| aTUNm63-Gibraltar-1755CE | N 36.28 W 6.09 | Conil | 1755 CE |
| aTUNm64-Gibraltar-1755CE | N 36.28 W 6.09 | Conil | 1755 CE |
| aTUNm65-Marseille-1800CE | N 43.30 E 5.37 | Marseille | 1800 CE |
| aTUNm66-Marseille-1800CE | N 43.30 E 5.37 | Marseille | 1800 CE |
| aTUNm67-Marseille-1800CE | N 43.30 E 5.37 | Marseille | 1800 CE |
| aTUNm68-Marseille-1800CE | N 43.30 E 5.37 | Marseille | 1800 CE |
| aTUNm69-Marseille-1800CE | N 43.30 E 5.37 | Marseille | 1800 CE |
| aTUNm70-Marseille-1800CE | N 43.30 E 5.37 | Marseille | 1800 CE |
| aTUNm71-Marseille-1800CE | N 43.30 E 5.37 | Marseille | 1800 CE |
| aTUNm72-Marseille-1800CE | N 43.30 E 5.37 | Marseille | 1800 CE |
| aTUNm73-Gibraltar-100CE | N 36.53 W 6.30 | Cadiz | 100 CE |
| aTUNm74-Gibraltar-100CE | N 36.53 W 6.30 | Cadiz | 100 CE |
| aTUNm75-Gibraltar-100CE | N 36.53 W 6.30 | Cadiz | 100 CE |
| aTUNm76-Gibraltar-100CE | N 36.53 W 6.30 | Cadiz | 100 CE |
| aTUNm77-Gibraltar-100CE | N 36.53 W 6.30 | Cadiz | 100 CE |
| aTUNm78-Sardinia-1500-1700CE | N 40.86 E 8.62 | Sassari | 1500-1700 CE |
| aTUNm79-Sardinia-1500-1700CE | N 40.86 E 8.62 | Sassari | 1500-1700 CE |
| aTUNm80-Sardinia-1500-1700CE | N 40.86 E 8.62 | Sassari | 1500-1700 CE |
| aTUNm81-Sardinia-1500-1700CE | N 40.86 E 8.62 | Sassari | 1500-1700 CE |
| aTUNm82-Sardinia-1500-1700CE | N 40.86 E 8.62 | Sassari | 1500-1700 CE |
| aTUNm83-Sardinia-1500-1700CE | N 40.86 E 8.62 | Sassari | 1500-1700 CE |
| aTUNm84-Sardinia-1500-1700CE | N 40.86 E 8.62 | Sassari | 1500-1700 CE |
| aTUNm85-Sardinia-1500-1700CE | N 40.86 E 8.62 | Sassari | 1500-1700 CE |
| aTUNm86-Sardinia-1500-1700CE | N 40.86 E 8.62 | Sassari | 1500-1700 CE |
| aTUNm87-Sardinia-1500-1700CE | N 40.86 E 8.62 | Sassari | 1500-1700 CE |
| aTUNm88-Sardinia-1500-1700CE | N 40.86 E 8.62 | Sassari | 1500-1700 CE |
| aTUNm89-Sardinia-1500-1700CE | N 40.86 E 8.62 | Sassari | 1500-1700 CE |
| aTUNm90-Sardinia-1500-1700CE | N 40.86 E 8.62 | Sassari | 1500-1700 CE |
| aTUNm91-Sardinia-1500-1700CE | N 40.86 E 8.62 | Sassari | 1500-1700 CE |
| aTUNm92-Sardinia-1500-1700CE | N 40.86 E 8.62 | Sassari | 1500-1700 CE |

**Table S3:**
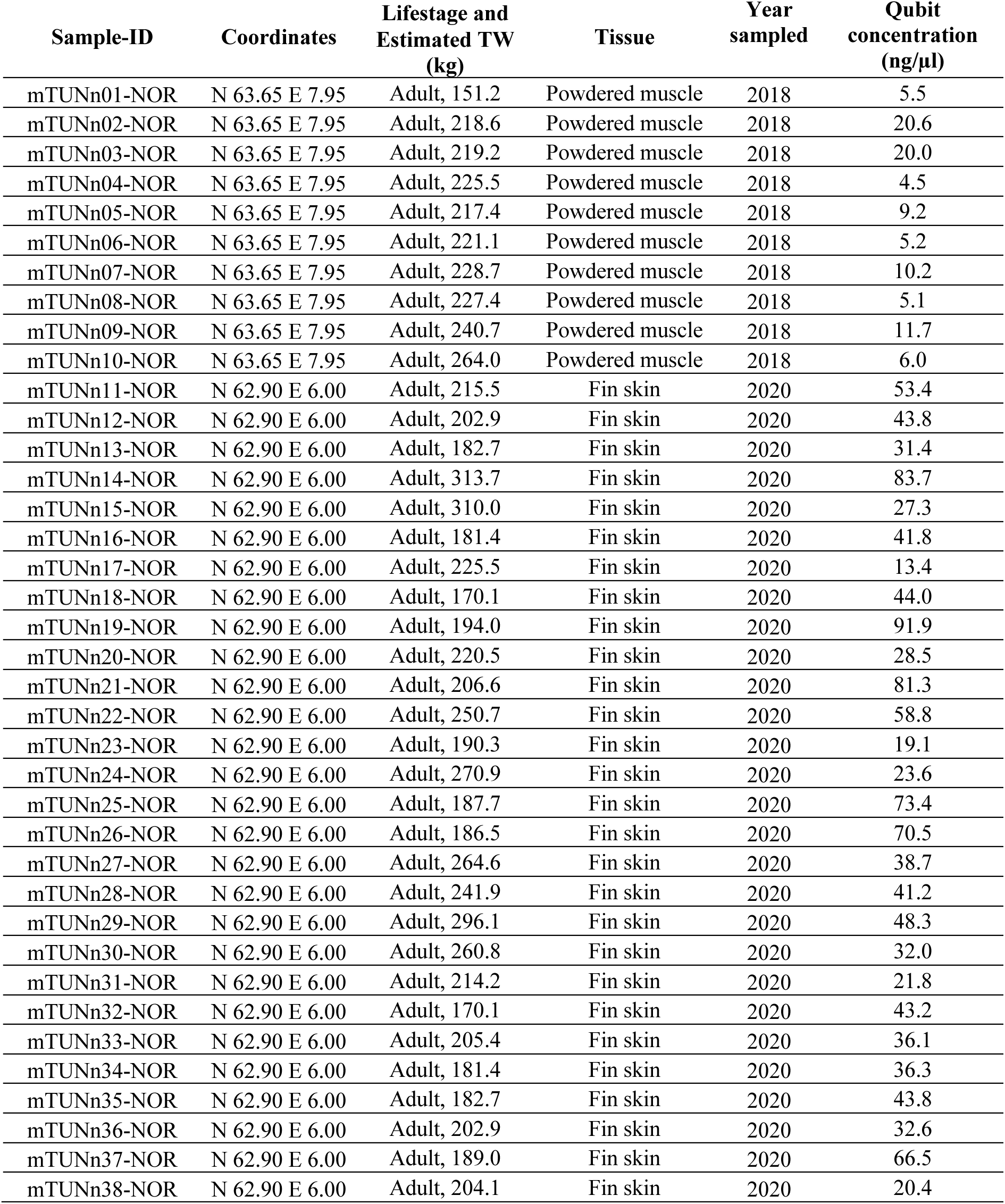
Sample locations, lifestage, tissue, date sampled and dsDNA concentration for the modern Atlantic bluefin samples from Norway.

| Sample-ID | Coordinates | Lifestage and<br>Estimated TW<br>(kg) | Tissue | Year<br>sampled | Qubit<br>concentration<br>(ng/μl) |
| --- | --- | --- | --- | --- | --- |
| mTUNn01-NOR | N 63.65 E 7.95 | Adult, 151.2 | Powdered muscle | 2018 | 5.5 |
| mTUNn02-NOR | N 63.65 E 7.95 | Adult, 218.6 | Powdered muscle | 2018 | 20.6 |
| mTUNn03-NOR | N 63.65 E 7.95 | Adult, 219.2 | Powdered muscle | 2018 | 20.0 |
| mTUNn04-NOR | N 63.65 E 7.95 | Adult, 225.5 | Powdered muscle | 2018 | 4.5 |
| mTUNn05-NOR | N 63.65 E 7.95 | Adult, 217.4 | Powdered muscle | 2018 | 9.2 |
| mTUNn06-NOR | N 63.65 E 7.95 | Adult, 221.1 | Powdered muscle | 2018 | 5.2 |
| mTUNn07-NOR | N 63.65 E 7.95 | Adult, 228.7 | Powdered muscle | 2018 | 10.2 |
| mTUNn08-NOR | N 63.65 E 7.95 | Adult, 227.4 | Powdered muscle | 2018 | 5.1 |
| mTUNn09-NOR | N 63.65 E 7.95 | Adult, 240.7 | Powdered muscle | 2018 | 11.7 |
| mTUNn10-NOR | N 63.65 E 7.95 | Adult, 264.0 | Powdered muscle | 2018 | 6.0 |
| mTUNn11-NOR | N 62.90 E 6.00 | Adult, 215.5 | Fin skin | 2020 | 53.4 |
| mTUNn12-NOR | N 62.90 E 6.00 | Adult, 202.9 | Fin skin | 2020 | 43.8 |
| mTUNn13-NOR | N 62.90 E 6.00 | Adult, 182.7 | Fin skin | 2020 | 31.4 |
| mTUNn14-NOR | N 62.90 E 6.00 | Adult, 313.7 | Fin skin | 2020 | 83.7 |
| mTUNn15-NOR | N 62.90 E 6.00 | Adult, 310.0 | Fin skin | 2020 | 27.3 |
| mTUNn16-NOR | N 62.90 E 6.00 | Adult, 181.4 | Fin skin | 2020 | 41.8 |
| mTUNn17-NOR | N 62.90 E 6.00 | Adult, 225.5 | Fin skin | 2020 | 13.4 |
| mTUNn18-NOR | N 62.90 E 6.00 | Adult, 170.1 | Fin skin | 2020 | 44.0 |
| mTUNn19-NOR | N 62.90 E 6.00 | Adult, 194.0 | Fin skin | 2020 | 91.9 |
| mTUNn20-NOR | N 62.90 E 6.00 | Adult, 220.5 | Fin skin | 2020 | 28.5 |
| mTUNn21-NOR | N 62.90 E 6.00 | Adult, 206.6 | Fin skin | 2020 | 81.3 |
| mTUNn22-NOR | N 62.90 E 6.00 | Adult, 250.7 | Fin skin | 2020 | 58.8 |
| mTUNn23-NOR | N 62.90 E 6.00 | Adult, 190.3 | Fin skin | 2020 | 19.1 |
| mTUNn24-NOR | N 62.90 E 6.00 | Adult, 270.9 | Fin skin | 2020 | 23.6 |
| mTUNn25-NOR | N 62.90 E 6.00 | Adult, 187.7 | Fin skin | 2020 | 73.4 |
| mTUNn26-NOR | N 62.90 E 6.00 | Adult, 186.5 | Fin skin | 2020 | 70.5 |
| mTUNn27-NOR | N 62.90 E 6.00 | Adult, 264.6 | Fin skin | 2020 | 38.7 |
| mTUNn28-NOR | N 62.90 E 6.00 | Adult, 241.9 | Fin skin | 2020 | 41.2 |
| mTUNn29-NOR | N 62.90 E 6.00 | Adult, 296.1 | Fin skin | 2020 | 48.3 |
| mTUNn30-NOR | N 62.90 E 6.00 | Adult, 260.8 | Fin skin | 2020 | 32.0 |
| mTUNn31-NOR | N 62.90 E 6.00 | Adult, 214.2 | Fin skin | 2020 | 21.8 |
| mTUNn32-NOR | N 62.90 E 6.00 | Adult, 170.1 | Fin skin | 2020 | 43.2 |
| mTUNn33-NOR | N 62.90 E 6.00 | Adult, 205.4 | Fin skin | 2020 | 36.1 |
| mTUNn34-NOR | N 62.90 E 6.00 | Adult, 181.4 | Fin skin | 2020 | 36.3 |
| mTUNn35-NOR | N 62.90 E 6.00 | Adult, 182.7 | Fin skin | 2020 | 43.8 |
| mTUNn36-NOR | N 62.90 E 6.00 | Adult, 202.9 | Fin skin | 2020 | 32.6 |
| mTUNn37-NOR | N 62.90 E 6.00 | Adult, 189.0 | Fin skin | 2020 | 66.5 |
| mTUNn38-NOR | N 62.90 E 6.00 | Adult, 204.1 | Fin skin | 2020 | 20.4 |

**Table S4:** Sample locations, lifestage, tissue and date sampled for the modern Atlantic bluefin samples from the Mediterranean and the Gulf of Mexico.

| Sample-ID | Coordinates | Location | Lifestage | Year sampled |
| --- | --- | --- | --- | --- |
| mTUNm01-EMED | N 35.51 E 33.42 | Cyprus | YoY | 2013 |
| mTUNm02-EMED | N 35.51 E 33.42 | Cyprus | YoY | 2013 |
| mTUNm03-EMED | N 35.51 E 33.42 | Cyprus | YoY | 2013 |
| mTUNm04-EMED | N 35.51 E 33.42 | Cyprus | YoY | 2013 |
| mTUNm05-EMED | N 35.51 E 33.42 | Cyprus | YoY | 2013 |
| mTUNm06-EMED | N 35.51 E 33.42 | Cyprus | YoY | 2013 |
| mTUNm07-EMED | N 35.51 E 33.42 | Cyprus | YoY | 2013 |
| mTUNm08-EMED | N 35.51 E 33.42 | Cyprus | YoY | 2013 |
| mTUNm09-EMED | N 35.51 E 33.42 | Cyprus | YoY | 2013 |
| mTUNm10-EMED | N 35.51 E 33.42 | Cyprus | YoY | 2013 |
| mTUNm11-WMED | N 39.27 E 2.07 | Palma | YoY | 2013 |
| mTUNm12-WMED | N 39.27 E 2.07 | Palma | YoY | 2013 |
| mTUNm13-WMED | N 39.27 E 2.07 | Palma | YoY | 2013 |
| mTUNm14-WMED | N 39.27 E 2.07 | Palma | YoY | 2013 |
| mTUNm15-WMED | N 39.27 E 2.07 | Palma | YoY | 2013 |
| mTUNm16-WMED | N 39.27 E 2.07 | Palma | YoY | 2013 |
| mTUNm17-WMED | N 39.27 E 2.07 | Palma | YoY | 2013 |
| mTUNm18-WMED | N 39.27 E 2.07 | Palma | YoY | 2013 |
| mTUNm19-WMED | N 39.27 E 2.07 | Palma | YoY | 2013 |
| mTUNm20-WMED | N 39.27 E 2.07 | Palma | YoY | 2013 |
| mTUNm31-CMED | N 36.93 E 13.18 | Sicily | YoY | 2013 |
| mTUNm32-CMED | N 36.93 E 13.18 | Sicily | YoY | 2013 |
| mTUNm33-CMED | N 36.93 E 13.18 | Sicily | YoY | 2013 |
| mTUNm34-CMED | N 36.93 E 13.18 | Sicily | YoY | 2013 |
| mTUNm35-CMED | N 36.93 E 13.18 | Sicily | YoY | 2013 |
| mTUNm36-CMED | N 36.93 E 13.18 | Sicily | YoY | 2013 |
| mTUNm37-CMED | N 36.93 E 13.18 | Sicily | YoY | 2013 |
| mTUNm38-CMED | N 36.93 E 13.18 | Sicily | YoY | 2013 |
| mTUNm39-CMED | N 36.93 E 13.18 | Sicily | YoY | 2013 |
| mTUNm40-CMED | N 36.93 E 13.18 | Sicily | YoY | 2013 |
| mTUNm41-GOM | N 26.12 W 87.78 | Gulf of Mexico | larvae | 2017 |
| mTUNm42-GOM | N 25.84 W 88.13 | Gulf of Mexico | larvae | 2017 |
| mTUNm43-GOM | N 28.33 W 87.25 | Gulf of Mexico | larvae | 2018 |
| mTUNm44-GOM | N 28.33 W 87.25 | Gulf of Mexico | larvae | 2018 |
| mTUNm45-GOM | N 28.33 W 87.25 | Gulf of Mexico | larvae | 2018 |
| mTUNm46-GOM | N 26.53 W 93.58 | Gulf of Mexico | larvae | 2014 |
| mTUNm47-GOM | N 26.53 W 93.58 | Gulf of Mexico | larvae | 2014 |
| mTUNm48-GOM | N 27.04 W 93.00 | Gulf of Mexico | larvae | 2014 |
| mTUNm49-GOM | N 28.00 W 87.76 | Gulf of Mexico | larvae | 2014 |
| mTUNm50-GOM | N 28.00 W 87.76 | Gulf of Mexico | larvae | 2014 |

**Table S5:** Sample locations, lifestage, tissue and date sampled for the modern albacore samples from the Bay of Biscay.

| Sample-ID | Coordinates | Location | Lifestage | Year sampled |
| --- | --- | --- | --- | --- |
| ALB-TUN-01 | N 44.46 W 3.12 | Bay of Biscay | Juvenile (5-15 kg) | 2010 |
| ALB-TUN-02 | N 44.46 W 3.12 | Bay of Biscay | Juvenile (5-15 kg) | 2010 |
| ALB-TUN-03 | N 44.46 W 3.12 | Bay of Biscay | Juvenile (5-15 kg) | 2010 |
| ALB-TUN-04 | N 44.46 W 3.12 | Bay of Biscay | Juvenile (5-15 kg) | 2010 |
| ALB-TUN-05 | N 44.46 W 3.12 | Bay of Biscay | Juvenile (5-15 kg) | 2010 |
| ALB-TUN-06 | N 44.46 W 3.12 | Bay of Biscay | Juvenile (5-15 kg) | 2010 |

**Table S6:** Pacific bluefin whole genome raw sequences downloaded from the DDBJ database (Kodama et al., 2012).

| Original sample-ID | Sample-ID | DDBJ identifier | Filename |
| --- | --- | --- | --- |
| PAC-DRR177383 | PBFT-TUN-01 | DRA008331 | DRR177383_1.fastq.bz2 |
|  |  |  | DRR177383_2.fastq.bz2 |
| PAC-DRR177395 | PBFT-TUN-02 | DRA008331 | DRR177395_1.fastq.bz2 |
|  |  |  | DRR177395_2.fastq.bz2 |
| PAC-DRR177400 | PBFT-TUN-03 | DRA008331 | DRR177400_1.fastq.bz2 |
|  |  |  | DRR177400_2.fastq.bz2 |
| PAC-DRR177401 | PBFT-TUN-04 | DRA008331 | DRR177401_1.fastq.bz2 |
|  |  |  | DRR177401_2.fastq.bz2 |
| PAC-DRR177402 | PBFT-TUN-05 | DRA008331 | DRR177402_1.fastq.bz2 |
|  |  |  | DRR177402_2.fastq.bz2 |
| PAC-DRR177403 | PBFT-TUN-06 | DRA008331 | DRR177403_1.fastq.bz2 |
|  |  |  | DRR177403_2.fastq.bz2 |
| PAC-DRR177404 | PBFT-TUN-07 | DRA008331 | DRR177404_1.fastq.bz2 |
|  |  |  | DRR177404_2.fastq.bz2 |
| PAC-DRR177405 | PBFT-TUN-08 | DRA008331 | DRR177405_1.fastq.bz2 |
|  |  |  | DRR177405_2.fastq.bz2 |
| PAC-DRR177406 | PBFT-TUN-09 | DRA008331 | DRR177406_1.fastq.bz2 |
|  |  |  | DRR177406_2.fastq.bz2 |

URL: https://ddbj.nig.ac.jp/public/ddbj_database/dra/fastq/DRA008/DRA008331/DRX167946/

**Table S7:**
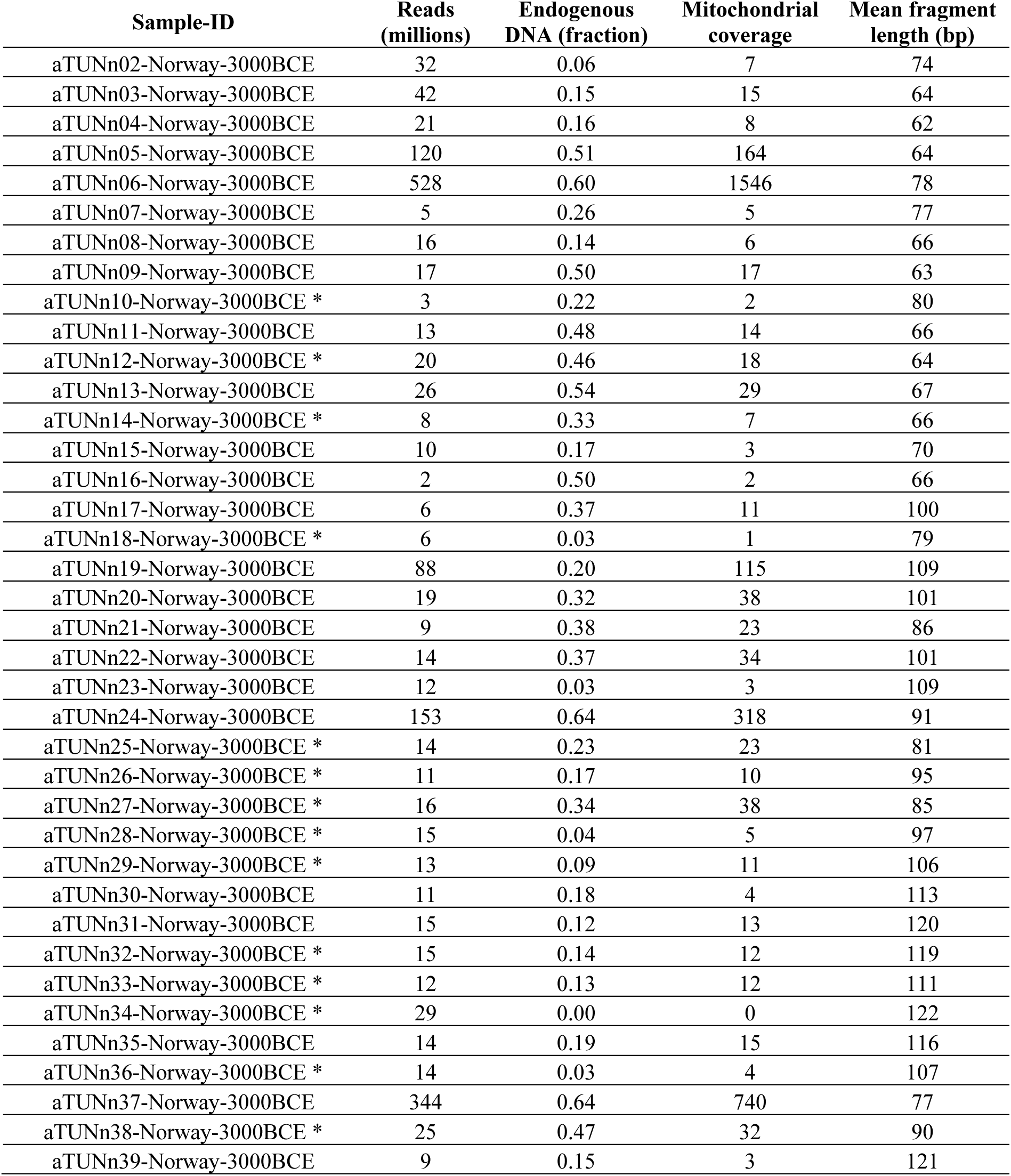
Summary statistics from Paleomix for the ancient Atlantic bluefin specimens from Norway. The endogenous content is calculated from the alignment to the Atlantic bluefin nuclear reference genome. Samples that were removed from further analyses are marked with a star (*).

| Sample-ID | Reads<br>(millions) | Endogenous<br>DNA (fraction) | Mitochondrial<br>coverage | Mean fragment<br>length (bp) |
| --- | --- | --- | --- | --- |
| aTUNn02-Norway-3000BCE | 32 | 0.06 | 7 | 74 |
| aTUNn03-Norway-3000BCE | 42 | 0.15 | 15 | 64 |
| aTUNn04-Norway-3000BCE | 21 | 0.16 | 8 | 62 |
| aTUNn05-Norway-3000BCE | 120 | 0.51 | 164 | 64 |
| aTUNn06-Norway-3000BCE | 528 | 0.60 | 1546 | 78 |
| aTUNn07-Norway-3000BCE | 5 | 0.26 | 5 | 77 |
| aTUNn08-Norway-3000BCE | 16 | 0.14 | 6 | 66 |
| aTUNn09-Norway-3000BCE | 17 | 0.50 | 17 | 63 |
| aTUNn10-Norway-3000BCE * | 3 | 0.22 | 2 | 80 |
| aTUNn11-Norway-3000BCE | 13 | 0.48 | 14 | 66 |
| aTUNn12-Norway-3000BCE * | 20 | 0.46 | 18 | 64 |
| aTUNn13-Norway-3000BCE | 26 | 0.54 | 29 | 67 |
| aTUNn14-Norway-3000BCE * | 8 | 0.33 | 7 | 66 |
| aTUNn15-Norway-3000BCE | 10 | 0.17 | 3 | 70 |
| aTUNn16-Norway-3000BCE | 2 | 0.50 | 2 | 66 |
| aTUNn17-Norway-3000BCE | 6 | 0.37 | 11 | 100 |
| aTUNn18-Norway-3000BCE * | 6 | 0.03 | 1 | 79 |
| aTUNn19-Norway-3000BCE | 88 | 0.20 | 115 | 109 |
| aTUNn20-Norway-3000BCE | 19 | 0.32 | 38 | 101 |
| aTUNn21-Norway-3000BCE | 9 | 0.38 | 23 | 86 |
| aTUNn22-Norway-3000BCE | 14 | 0.37 | 34 | 101 |
| aTUNn23-Norway-3000BCE | 12 | 0.03 | 3 | 109 |
| aTUNn24-Norway-3000BCE | 153 | 0.64 | 318 | 91 |
| aTUNn25-Norway-3000BCE * | 14 | 0.23 | 23 | 81 |
| aTUNn26-Norway-3000BCE * | 11 | 0.17 | 10 | 95 |
| aTUNn27-Norway-3000BCE * | 16 | 0.34 | 38 | 85 |
| aTUNn28-Norway-3000BCE * | 15 | 0.04 | 5 | 97 |
| aTUNn29-Norway-3000BCE * | 13 | 0.09 | 11 | 106 |
| aTUNn30-Norway-3000BCE | 11 | 0.18 | 4 | 113 |
| aTUNn31-Norway-3000BCE | 15 | 0.12 | 13 | 120 |
| aTUNn32-Norway-3000BCE * | 15 | 0.14 | 12 | 119 |
| aTUNn33-Norway-3000BCE * | 12 | 0.13 | 12 | 111 |
| aTUNn34-Norway-3000BCE * | 29 | 0.00 | 0 | 122 |
| aTUNn35-Norway-3000BCE | 14 | 0.19 | 15 | 116 |
| aTUNn36-Norway-3000BCE * | 14 | 0.03 | 4 | 107 |
| aTUNn37-Norway-3000BCE | 344 | 0.64 | 740 | 77 |
| aTUNn38-Norway-3000BCE * | 25 | 0.47 | 32 | 90 |
| aTUNn39-Norway-3000BCE | 9 | 0.15 | 3 | 121 |

**Table S8:** Summary statistics from Paleomix for the ancient Atlantic bluefin specimens from the Mediterranean. The endogenous content is calculated from the alignment to the Atlantic bluefin nuclear reference genome. Samples that were removed from further analyses are marked with a star (*).

| Sample-ID | Reads<br>(millions) | Endogenous<br>DNA (fraction) | Mitochondrial<br>coverage | Mean fragment<br>length (bp) |
| --- | --- | --- | --- | --- |
| aTUNm01-Sicily-900-1200CE | 16 | 0.07 | 2 | 83 |
| aTUNm02-Sicily-900-1200CE | 5 | 0.17 | 5 | 79 |
| aTUNm03-Sicily-900-1200CE * | 8 | 0.19 | 5 | 68 |
| aTUNm04-Sicily-900-1200CE * | 5 | 0.17 | 3 | 86 |
| aTUNm05-Sicily-900-1200CE | 18 | 0.12 | 10 | 91 |
| aTUNm06-Sicily-900-1200CE | 6 | 0.21 | 6 | 92 |
| aTUNm07-Sicily-900-1200CE | 6 | 0.52 | 16 | 86 |
| aTUNm08-Sicily-900-1200CE | 5 | 0.22 | 4 | 103 |
| aTUNm09-Sicily-900-1200CE | 8 | 0.07 | 2 | 93 |
| aTUNm10-Sicily-900-1200CE | 16 | 0.58 | 73 | 73 |
| aTUNm11-Sicily-900-1200CE | 7 | 0.59 | 25 | 79 |
| aTUNm12-Sicily-900-1200CE | 7 | 0.12 | 4 | 92 |
| aTUNm13-Zliten-1925CE | 7 | 0.38 | 12 | 70 |
| aTUNm14-Zliten-1925CE | 8 | 0.36 | 15 | 77 |
| aTUNm15-Zliten-1925CE | 7 | 0.06 | 3 | 80 |
| aTUNm16-Zliten-1925CE | 6 | 0.11 | 4 | 68 |
| aTUNm17-Zliten-1925CE | 6 | 0.33 | 5 | 69 |
| aTUNm18-Zliten-1925CE | 8 | 0.40 | 12 | 72 |
| aTUNm19-Zliten-1925CE | 7 | 0.25 | 6 | 75 |
| aTUNm20-Zliten-1925CE | 4 | 0.46 | 7 | 68 |
| aTUNm21-Zliten-1925CE * | 6 | 0.06 | 3 | 70 |
| aTUNm22-Zliten-1925CE | 8 | 0.37 | 13 | 67 |
| aTUNm23-Zliten-1925CE | 7 | 0.15 | 5 | 63 |
| aTUNm24-Zliten-1925CE | 7 | 0.17 | 4 | 66 |
| aTUNm25-Zliten-1925CE | 6 | 0.45 | 9 | 69 |
| aTUNm26-Zliten-1925CE * | 9 | 0.08 | 4 | 72 |
| aTUNm27-Zliten-1925CE | 8 | 0.07 | 5 | 83 |
| aTUNm28-Zliten-1925CE | 9 | 0.25 | 9 | 77 |
| aTUNm29-Zliten-1925CE | 9 | 0.18 | 6 | 77 |
| aTUNm30-Zliten-1925CE | 10 | 0.04 | 3 | 86 |
| aTUNm31-Zliten-1925CE | 7 | 0.49 | 10 | 80 |
| aTUNm32-Zliten-1925CE | 9 | 0.28 | 17 | 83 |
| aTUNm33-Zliten-1925CE | 7 | 0.26 | 6 | 74 |
| aTUNm34-Zliten-1925CE | 6 | 0.10 | 4 | 80 |
| aTUNm35-Istanbul-1941CE | 6 | 0.11 | 1 | 111 |
| aTUNm36-Istanbul-1941CE | 10 | 0.27 | 10 | 72 |
| aTUNm37-Istanbul-800-1200CE | 5 | 0.51 | 22 | 77 |
| aTUNm38-Istanbul-800-1200CE | 5 | 0.47 | 9 | 70 |
| aTUNm39-Istanbul-800-1200CE | 5 | 0.24 | 7 | 82 |
| aTUNm40-Istanbul-800-1200CE * | 4 | 0.45 | 19 | 91 |
| aTUNm41-Istanbul-800-1200CE | 5 | 0.57 | 25 | 76 |
| aTUNm42-Istanbul-800-1200CE | 4 | 0.36 | 7 | 90 |
| aTUNm43-Istanbul-800-1200CE | 4 | 0.32 | 4 | 74 |
| aTUNm44-Istanbul-800-1200CE | 5 | 0.16 | 3 | 76 |
| aTUNm45-Istanbul-800-1200CE | 4 | 0.59 | 16 | 75 |
| aTUNm46-Istanbul-800-1200CE | 4 | 0.53 | 18 | 83 |
| aTUNm47-Istanbul-800-1200CE | 4 | 0.61 | 12 | 77 |
| aTUNm48-Istanbul-800-1200CE * | 4 | 0.09 | 2 | 78 |
| aTUNm49-Istanbul-800-1200CE | 4 | 0.22 | 5 | 79 |
| aTUNm50-Istanbul-800-1200CE | 6 | 0.50 | 12 | 82 |
| aTUNm51-Istanbul-800-1200CE | 10 | 0.46 | 31 | 75 |
| aTUNm52-Istanbul-800-1200CE | 14 | 0.39 | 64 | 78 |
| aTUNm53-Istanbul-800-1200CE | 6 | 0.38 | 8 | 76 |
| aTUNm54-Istanbul-800-1200CE | 9 | 0.33 | 15 | 87 |
| aTUNm55-Istanbul-800-1200CE | 6 | 0.42 | 10 | 77 |
| aTUNm56-Istanbul-800-1200CE | 10 | 0.47 | 17 | 77 |
| aTUNm57-Istanbul-800-1200CE | 4 | 0.55 | 25 | 96 |
| aTUNm58-Gibraltar-1755CE | 10 | 0.06 | 2 | 82 |
| aTUNm59-Gibraltar-1755CE | 8 | 0.10 | 3 | 79 |
| aTUNm60-Gibraltar-1755CE | 6 | 0.10 | 3 | 87 |
| aTUNm61-Gibraltar-1755CE | 8 | 0.08 | 3 | 82 |
| aTUNm62-Gibraltar-1755CE | 6 | 0.16 | 2 | 85 |
| aTUNm63-Gibraltar-1755CE | 5 | 0.10 | 3 | 89 |
| aTUNm64-Gibraltar-1755CE | 14 | 0.06 | 3 | 83 |
| aTUNm65-Marseille-1800CE | 6 | 0.34 | 10 | 72 |
| aTUNm66-Marseille-1800CE | 7 | 0.35 | 69 | 95 |
| aTUNm67-Marseille-1800CE | 9 | 0.22 | 7 | 99 |
| aTUNm68-Marseille-1800CE | 10 | 0.24 | 6 | 82 |
| aTUNm69-Marseille-1800CE | 3 | 0.28 | 8 | 82 |
| aTUNm70-Marseille-1800CE | 6 | 0.25 | 12 | 82 |
| aTUNm71-Marseille-1800CE | 9 | 0.18 | 27 | 92 |
| aTUNm72-Marseille-1800CE | 38 | 0.12 | 14 | 72 |
| aTUNm73-Gibraltar-100CE | 22 | 0.05 | 11 | 92 |
| aTUNm74-Gibraltar-100CE * | 13 | 0.07 | 2 | 83 |
| aTUNm75-Gibraltar-100CE | 21 | 0.13 | 10 | 85 |
| aTUNm76-Gibraltar-100CE | 13 | 0.09 | 5 | 99 |
| aTUNm77-Gibraltar-100CE | 15 | 0.08 | 6 | 82 |
| aTUNm78-Sardinia-1500-1700CE | 5 | 0.39 | 8 | 103 |
| aTUNm79-Sardinia-1500-1700CE | 11 | 0.57 | 29 | 75 |
| aTUNm80-Sardinia-1500-1700CE | 6 | 0.06 | 4 | 105 |
| aTUNm81-Sardinia-1500-1700CE | 6 | 0.39 | 10 | 86 |
| aTUNm82-Sardinia-1500-1700CE | 5 | 0.40 | 15 | 93 |
| aTUNm83-Sardinia-1500-1700CE | 11 | 0.05 | 2 | 89 |
| aTUNm84-Sardinia-1500-1700CE | 5 | 0.39 | 14 | 90 |
| aTUNm85-Sardinia-1500-1700CE | 13 | 0.56 | 34 | 81 |
| aTUNm86-Sardinia-1500-1700CE | 7 | 0.14 | 5 | 99 |
| aTUNm87-Sardinia-1500-1700CE | 14 | 0.58 | 58 | 93 |
| aTUNm88-Sardinia-1500-1700CE | 9 | 0.49 | 13 | 84 |
| aTUNm89-Sardinia-1500-1700CE | 6 | 0.57 | 15 | 80 |
| aTUNm90-Sardinia-1500-1700CE | 10 | 0.59 | 31 | 87 |
| aTUNm91-Sardinia-1500-1700CE | 3 | 0.50 | 11 | 90 |
| aTUNm92-Sardinia-1500-1700CE | 3 | 0.36 | 7 | 96 |

**Table S9:** Summary statistics from Paleomix for the modern Atlantic bluefin specimens from Norway. The endogenous content is calculated from the alignment to the Atlantic bluefin nuclear reference genome.

| Sample-ID | Reads<br>(millions) | Endogenous<br>DNA (fraction) | Mitochondrial<br>coverage | Mean fragment<br>length (bp) |
| --- | --- | --- | --- | --- |
| mTUNn01-NOR | 75 | 0.69 | 2784 | 178 |
| mTUNn02-NOR | 59 | 0.73 | 1019 | 170 |
| mTUNn03-NOR | 56 | 0.72 | 1508 | 172 |
| mTUNn04-NOR | 79 | 0.75 | 1150 | 178 |
| mTUNn05-NOR | 69 | 0.76 | 823 | 172 |
| mTUNn06-NOR | 79 | 0.75 | 1633 | 169 |
| mTUNn07-NOR | 73 | 0.77 | 1163 | 170 |
| mTUNn08-NOR | 76 | 0.75 | 1811 | 175 |
| mTUNn09-NOR | 90 | 0.76 | 1960 | 171 |
| mTUNn10-NOR | 93 | 0.75 | 2345 | 171 |
| mTUNn11-NOR | 21 | 0.76 | 273 | 168 |
| mTUNn12-NOR | 19 | 0.75 | 307 | 168 |
| mTUNn13-NOR | 23 | 0.78 | 296 | 168 |
| mTUNn14-NOR | 19 | 0.76 | 247 | 170 |
| mTUNn15-NOR | 18 | 0.76 | 283 | 166 |
| mTUNn16-NOR | 22 | 0.76 | 370 | 170 |
| mTUNn17-NOR | 22 | 0.75 | 260 | 168 |
| mTUNn18-NOR | 33 | 0.75 | 512 | 169 |
| mTUNn19-NOR | 22 | 0.77 | 228 | 167 |
| mTUNn20-NOR | 19 | 0.75 | 251 | 168 |
| mTUNn21-NOR | 19 | 0.76 | 520 | 168 |
| mTUNn22-NOR | 17 | 0.77 | 286 | 166 |
| mTUNn23-NOR | 17 | 0.77 | 263 | 167 |
| mTUNn24-NOR | 23 | 0.77 | 301 | 168 |
| mTUNn25-NOR | 23 | 0.75 | 381 | 169 |
| mTUNn26-NOR | 21 | 0.76 | 389 | 169 |
| mTUNn27-NOR | 25 | 0.76 | 343 | 168 |
| mTUNn28-NOR | 25 | 0.76 | 349 | 166 |
| mTUNn29-NOR | 22 | 0.75 | 448 | 168 |
| mTUNn30-NOR | 23 | 0.77 | 419 | 169 |
| mTUNn31-NOR | 27 | 0.77 | 687 | 166 |
| mTUNn32-NOR | 22 | 0.78 | 543 | 168 |
| mTUNn33-NOR | 22 | 0.75 | 458 | 169 |
| mTUNn34-NOR | 18 | 0.76 | 417 | 166 |
| mTUNn35-NOR | 24 | 0.76 | 472 | 167 |
| mTUNn36-NOR | 23 | 0.76 | 356 | 168 |
| mTUNn37-NOR | 24 | 0.77 | 419 | 166 |
| mTUNn38-NOR | 24 | 0.75 | 425 | 169 |

**Table S10:** Summary statistics from Paleomix for the modern Atlantic bluefin specimens from the Mediterranean and the Gulf of Mexico. The endogenous content is calculated from the alignment to the Atlantic bluefin nuclear reference genome.

| Sample-ID | Reads<br>(millions) | Endogenous<br>DNA (fraction) | Mitochondrial<br>coverage | Mean fragment<br>length (bp) |
| --- | --- | --- | --- | --- |
| mTUNm01-EMED | 24 | 0.68 | 716 | 112 |
| mTUNm02-EMED | 25 | 0.68 | 704 | 124 |
| mTUNm03-EMED | 22 | 0.69 | 336 | 118 |
| mTUNm04-EMED | 13 | 0.80 | 445 | 156 |
| mTUNm05-EMED | 48 | 0.48 | 2089 | 120 |
| mTUNm06-EMED | 37 | 0.49 | 1600 | 115 |
| mTUNm07-EMED | 31 | 0.67 | 708 | 105 |
| mTUNm08-EMED | 21 | 0.72 | 230 | 102 |
| mTUNm09-EMED | 19 | 0.71 | 363 | 100 |
| mTUNm10-EMED | 23 | 0.73 | 538 | 103 |
| mTUNm11-WMED | 24 | 0.68 | 1021 | 83 |
| mTUNm12-WMED | 27 | 0.70 | 660 | 112 |
| mTUNm13-WMED | 14 | 0.59 | 574 | 104 |
| mTUNm14-WMED | 20 | 0.70 | 396 | 81 |
| mTUNm15-WMED | 26 | 0.66 | 840 | 119 |
| mTUNm16-WMED | 46 | 0.61 | 1462 | 80 |
| mTUNm17-WMED | 63 | 0.58 | 1212 | 67 |
| mTUNm18-WMED | 60 | 0.57 | 1183 | 62 |
| mTUNm19-WMED | 41 | 0.69 | 410 | 95 |
| mTUNm20-WMED | 74 | 0.65 | 460 | 70 |
| mTUNm31-CMED | 36 | 0.35 | 1180 | 122 |
| mTUNm32-CMED | 32 | 0.30 | 666 | 126 |
| mTUNm33-CMED | 54 | 0.60 | 1385 | 69 |
| mTUNm34-CMED | 56 | 0.34 | 976 | 128 |
| mTUNm35-CMED | 38 | 0.61 | 448 | 71 |
| mTUNm36-CMED | 32 | 0.71 | 745 | 106 |
| mTUNm37-CMED | 45 | 0.55 | 530 | 104 |
| mTUNm38-CMED | 36 | 0.35 | 176 | 128 |
| mTUNm39-CMED | 62 | 0.58 | 705 | 81 |
| mTUNm40-CMED | 51 | 0.58 | 533 | 60 |
| mTUNm41-GOM | 80 | 0.66 | 696 | 69 |
| mTUNm42-GOM | 51 | 0.38 | 972 | 111 |
| mTUNm43-GOM | 30 | 0.49 | 565 | 106 |
| mTUNm44-GOM | 52 | 0.56 | 881 | 100 |
| mTUNm45-GOM | 44 | 0.47 | 966 | 128 |
| mTUNm46-GOM | 26 | 0.69 | 319 | 78 |
| mTUNm47-GOM | 46 | 0.69 | 49 | 75 |
| mTUNm48-GOM | 26 | 0.65 | 34 | 67 |
| mTUNm49-GOM | 67 | 0.55 | 910 | 102 |
| mTUNm50-GOM | 53 | 0.60 | 430 | 66 |

**Table S11:** Summary statistics from Paleomix for the modern albacore specimens from the Bay of Biscay. The endogenous content is calculated from the alignment to the Atlantic bluefin nuclear reference genome.

| Sample-ID | Reads<br>(millions) | Endogenous<br>DNA (fraction) | Mitochondrial<br>coverage | Mean fragment<br>length (bp) |
| --- | --- | --- | --- | --- |
| ALB-TUN-01 | 37 | 0.59 | 277 | 99 |
| ALB-TUN-02 | 19 | 0.66 | 219 | 117 |
| ALB-TUN-03 | 28 | 0.65 | 103 | 124 |
| ALB-TUN-04 | 27 | 0.66 | 377 | 74 |
| ALB-TUN-05 | 12 | 0.70 | 243 | 95 |
| ALB-TUN-06 | 25 | 0.68 | 109 | 99 |

**Table S12:** Summary statistics from Paleomix for the modern Pacific bluefin specimens from the Nansei Islands. Whole genome raw sequence data downloaded from the DDBJ database (Kodama et al., 2012). The endogenous content is calculated from the alignment to the Atlantic bluefin nuclear reference genome.

| Sample-ID | Reads<br>(millions) | Endogenous<br>DNA (fraction) | Mitochondrial<br>coverage | Mean fragment<br>length (bp) |
| --- | --- | --- | --- | --- |
| PBFT-TUN-01 | 121 | 0.72 | 2657 | 152 |
| PBFT-TUN-02 | 127 | 0.74 | 5107 | 151 |
| PBFT-TUN-03 | 109 | 0.76 | 2346 | 151 |
| PBFT-TUN-04 | 112 | 0.73 | 1361 | 151 |
| PBFT-TUN-05 | 107 | 0.76 | 4005 | 151 |
| PBFT-TUN-06 | 112 | 0.75 | 3345 | 151 |
| PBFT-TUN-07 | 99 | 0.76 | 3061 | 151 |
| PBFT-TUN-08 | 126 | 0.75 | 3465 | 151 |
| PBFT-TUN-09 | 114 | 0.74 | 4555 | 151 |

**Table S13:** Introgressed individuals. Samples that were removed from further analyses due to being identical with other samples and/or having high missingness are marked with a star (*).

| **Sample-ID** | **Mitochondrial haplotype** |
| --- | --- |
| aTUNn21-Norway-3000BCE | Pacific bluefin |
| aTUNn22-Norway-3000BCE | Pacific bluefin |
| aTUNn25-Norway-3000BCE * | Pacific bluefin |
| aTUNn26-Norway-3000BCE * | Pacific bluefin |
| aTUNn27-Norway-3000BCE * | Pacific bluefin |
| aTUNn28-Norway-3000BCE * | Pacific bluefin |
| aTUNn29-Norway-3000BCE * | Pacific bluefin |
| aTUNn33-Norway-3000BCE * | Pacific bluefin |
| aTUNn34-Norway-3000BCE * | Pacific bluefin |
| aTUNn36-Norway-3000BCE * | Pacific bluefin |
| aTUNm02-Sicily-900-1200CE | Albacore |
| aTUNm04-Sicily-900-1200CE * | Albacore |
| aTUNm06-Sicily-900-1200CE | Albacore |
| aTUNm39-Istanbul-800-1200CE | Pacific bluefin |
| aTUNm46-Istanbul-800-1200CE | Albacore |
| aTUNm81-Sardinia-1500-1700CE | Pacific bluefin |
| mTUNn05-NOR | Albacore |
| mTUNn17-NOR | Pacific bluefin |
| mTUNn35-NOR | Pacific bluefin |
| mTUNm15-WMED | Albacore |

**Table S14:**
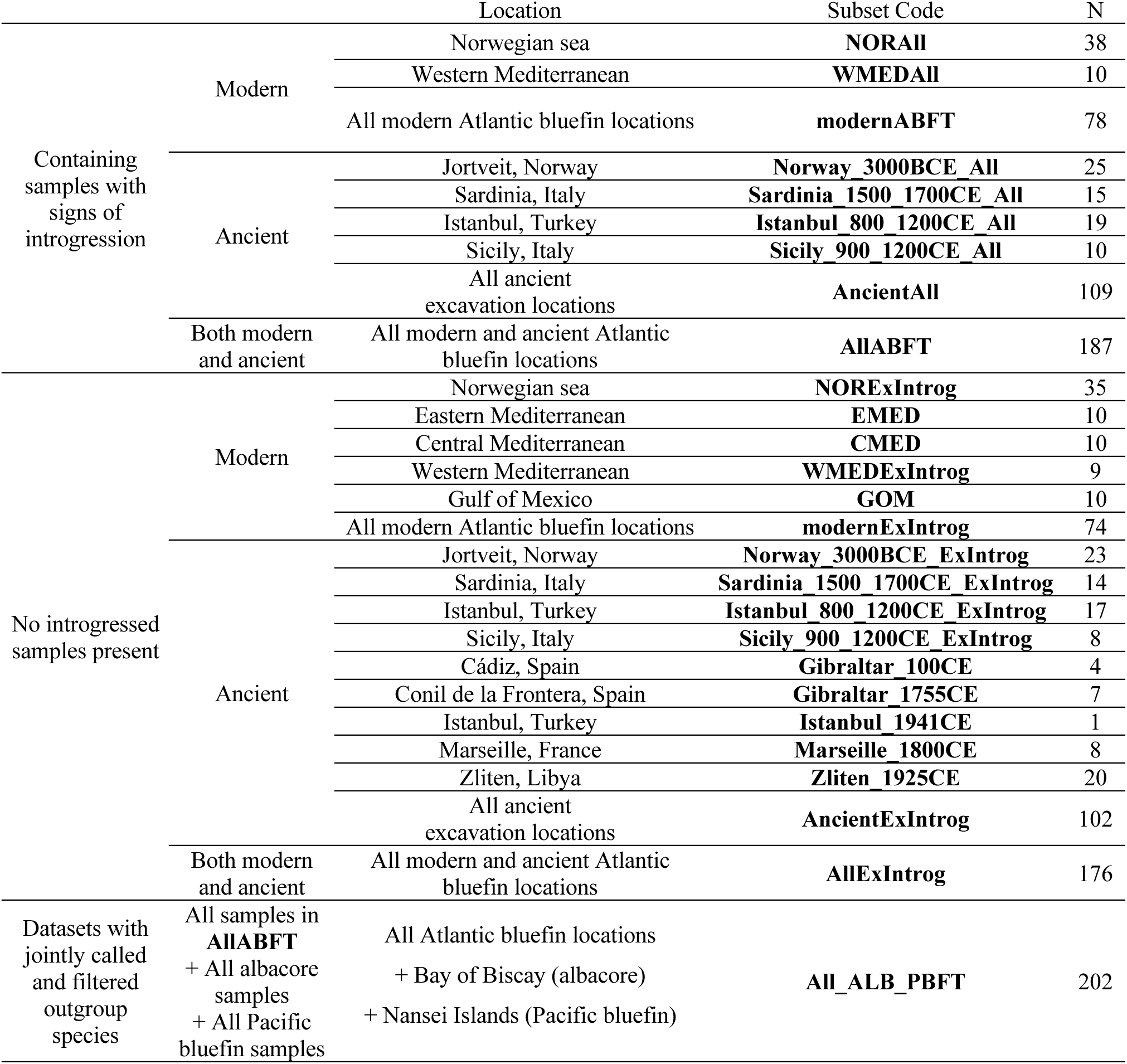
Jointly called and filtered datasets, used in population genomic analyses (N = number of samples in each dataset).

In the table above, the following datasets were used for population genomic analyses:

- AllABFT: Used for LociMissingness analyses (Figure S5) and calculation of ΦST and d_xy_ (Figure 3B). No outgroup.
- All_ALB_PBFT: Used for interspecific PCA (Figure 2A) with no outgroup. It was also used for haplotype network (Figure 2C) and ML and Bayesian phylogenies (Figure 2D and Figure S6) using one katsuwonus pelamis as outgroup (NCBI Reference Sequence: NC_005316.1)
- AllExIntrog: Used for intraspecific PCA (Figure S8) and calculation of ΦST and d_xy_ (Figure 3A) with no outgroup. It was also used for intraspecific haplotype network (Figure S9), as well as ML and Bayesian phylogenies (Figure S10) using one Pacific bluefin as outgroup (NCBI Reference Sequence: NC_008455.1)

All other datasets were only used for the calculation of population genomic statistics (Table S15).

**Table S15:** Population genomic statistics of the separately called and filtered datasets. N = number of samples, hD = haplotype diversity, Nh = number of haplotypes, S = number of segregating sites, π = nucleotide diversity, TD = Tajima’s D. Significance levels (**=0.01, *=0.05, n.s.=not significant) are indicated for the TD values. Datasets containing introgressed samples have been highlighted in beige color.

| Subset | N | hD | Nh | S | $\pi$ | TD | $\pi(\text{bs})$ | TD(bs) |
| --- | --- | --- | --- | --- | --- | --- | --- | --- |
| AllABFT_mt | 186 | 0.998 | 160 | 809 | 0.0033 | -3.23 (**) | 0.0027 | -2.49 |
| AllExIntrogt_mt | 175 | 0.999 | 158 | 475 | 0.0009 | -3.23 (**) | 0.0007 | -2.82 |
| modernABFT_mt | 78 | 0.998 | 72 | 699 | 0.0035 | -2.14 (*) | 0.0035 | -1.18 |
| modernExIntrogt_mt | 74 | 0.999 | 70 | 300 | 0.0012 | -2.49 (*) | 0.0011 | -1.20 |
| AncientAll_mt | 108 | 0.998 | 97 | 710 | 0.0034 | -3.37 (**) | 0.0024 | -3.42 |
| AncientExIntrogt_mt | 101 | 0.997 | 90 | 332 | 0.0008 | -3.37 (**) | 0.0005 | -3.45 |
| NORAll_mt | 38 | 0.997 | 36 | 621 | 0.0049 | -1.73 (n.s.) | 0.0050 | -0.92 |
| NORExIntrogt_mt | 35 | 0.998 | 34 | 187 | 0.0012 | -2.10 (*) | 0.0012 | -0.85 |
| WMEDAll_mt | 10 | 1.000 | 10 | 477 | 0.0060 | -2.14 (*) | 0.0060 | -1.28 |
| WMEDExIntrogt_mt | 9 | 1.000 | 9 | 74 | 0.0011 | -1.71 (n.s.) | 0.0011 | -1.21 |
| EMED_mt | 10 | 1.000 | 10 | 90 | 0.0013 | -1.59 (n.s.) | 0.0013 | -0.93 |
| CMED_mt | 10 | 1.000 | 10 | 90 | 0.0012 | -1.87 (n.s.) | 0.0012 | -1.21 |
| GOM_mt | 10 | 1.000 | 10 | 78 | 0.0010 | -2.17 (*) | 0.0010 | -1.58 |
| Norway_3000BCE_All | 24 | 1.000 | 24 | 556 | 0.0046 | -3.91 (**) | 0.0029 | -3.68 |
| Norway_3000BCE_ExIntrogt | 23 | 1.000 | 22 | 201 | 0.0012 | -3.92 (**) | 0.0008 | -3.65 |
| Sardinia_1500_1700CE_All | 15 | 1.000 | 15 | 504 | 0.0042 | -3.31 (**) | 0.0035 | -2.46 |
| Sardinia_1500_1700CE_ExIntrogt | 14 | 1.000 | 14 | 95 | 0.0009 | -3.47 (**) | 0.0007 | -2.65 |
| Istanbul_800_1200CE_All | 19 | 1.000 | 19 | 542 | 0.0054 | -2.90 (**) | 0.0049 | -1.94 |
| Istanbul_800_1200CE_ExIntrogt | 17 | 1.000 | 17 | 85 | 0.0009 | -3.04 (**) | 0.0007 | -2.01 |
| Sicily_900_1200CE_All | 10 | 1.000 | 10 | 411 | 0.0062 | -4.18 (**) | 0.0037 | -4.15 |
| Sicily_900_1200CE_ExIntrogt | 8 | 1.000 | 8 | 43 | 0.0006 | -4.55 (**) | 0.0003 | -4.64 |
| Gibraltar_100CE | 4 | 1.000 | 4 | 27 | 0.0008 | -0.63 (n.s.) | n.a. | n.a. |
| Gibraltar_1755CE | 7 | 1.000 | 7 | 30 | 0.0003 | -5.17(**) | 0.0001 | -6.27 |
| Istanbul_1941CE | 1 | n.a. | 1 | 0 | n.a. | n.a. | n.a. | n.a. |
| Marseille_1800CE | 8 | 1.000 | 8 | 40 | 0.0006 | -2.96 (**) | 0.0005 | -2.70 |
| Zliten_1925CE | 20 | 1.000 | 20 | 111 | 0.0008 | -3.75 (**) | 0.0006 | -3.38 |

### Section 3: Supplementary Figures

**Figure S1:**
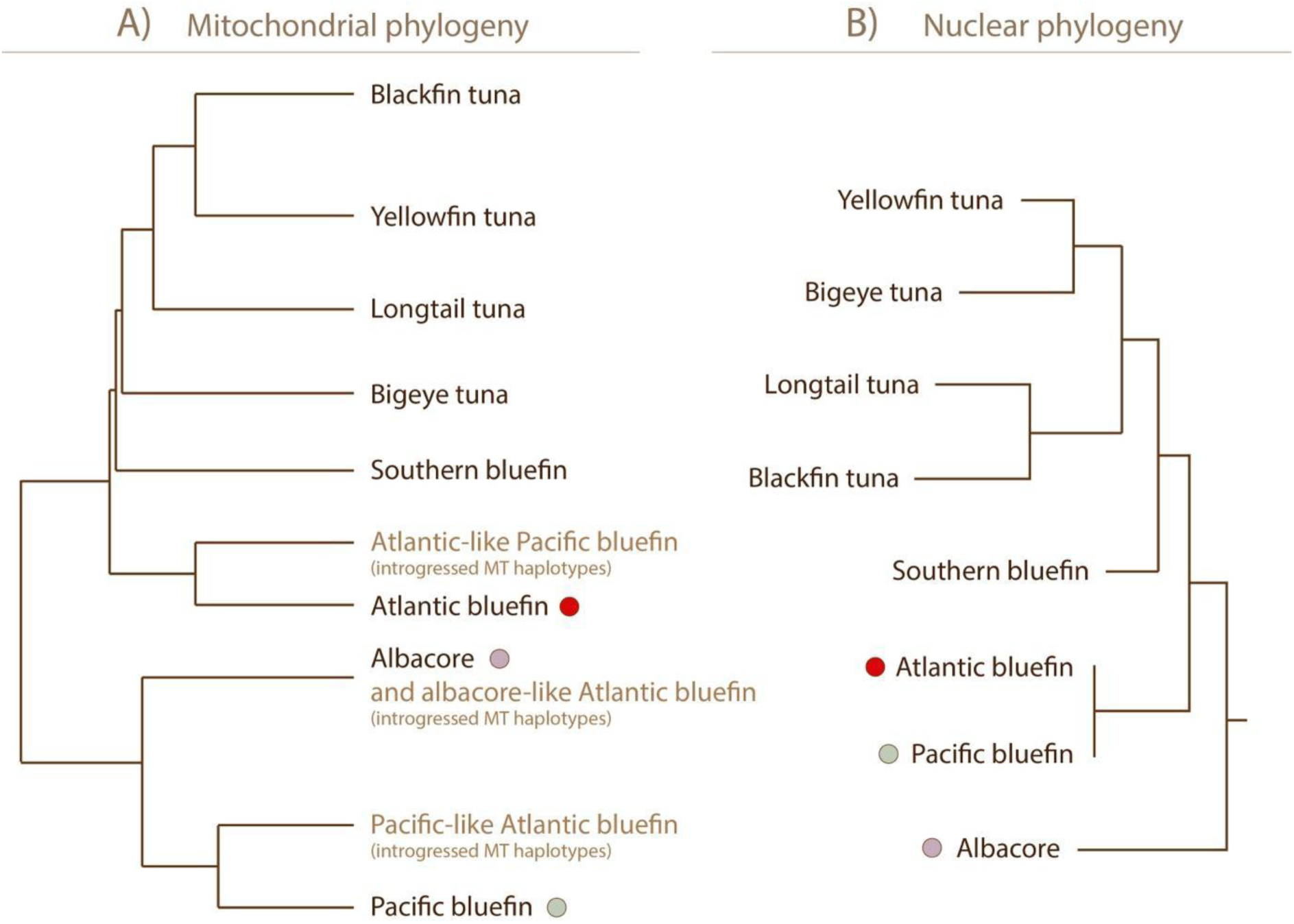
Thunnus phylogenies based on A) the mitochondrial control region adapted from Viñas & Tudela (2009) and B) genome-wide nuclear markers adapted from Díaz-Arce et al. (2016). Tip labels in light brown represent introgressed mitochondrial sequences. The colored circles mark species presented in this study.

**Figure S2:**
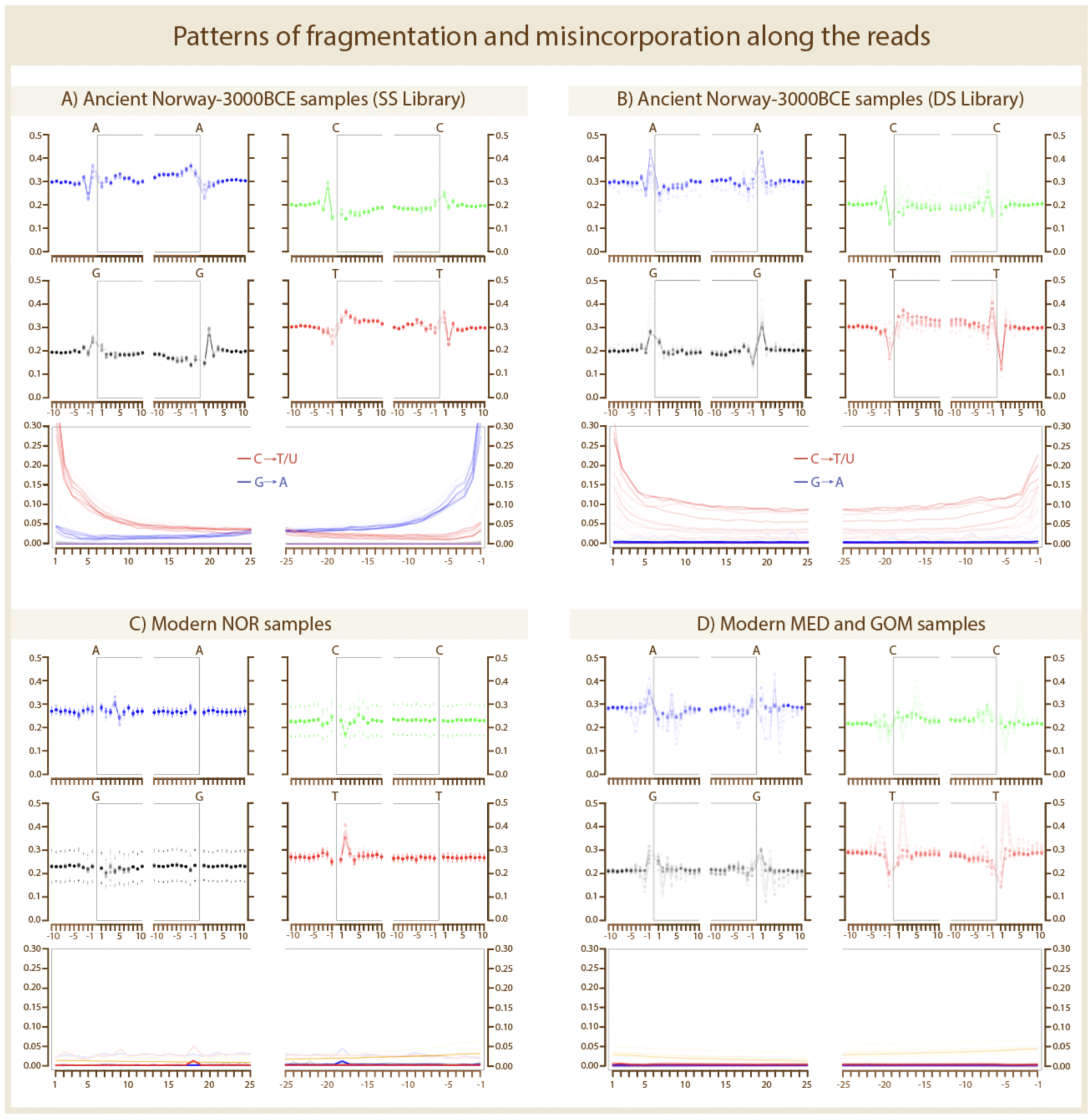
Fragmentation (upper panels) and misincorporation (lower panels) plots from mapDamage v.2.0.9. The fragmentation plots show the base frequency inside and surrounding the read, where the grey box indicates the location of where the reads have mapped to the reference. The misincorporation plots show the rate of substitutions along the positions of the read ends, relative to the reference (Red: C to T. Blue: G to A. Grey: All other substitutions. Green: Deletions. Purple: Insertions. Orange: Soft-clipped bases).

**Figure S3:**
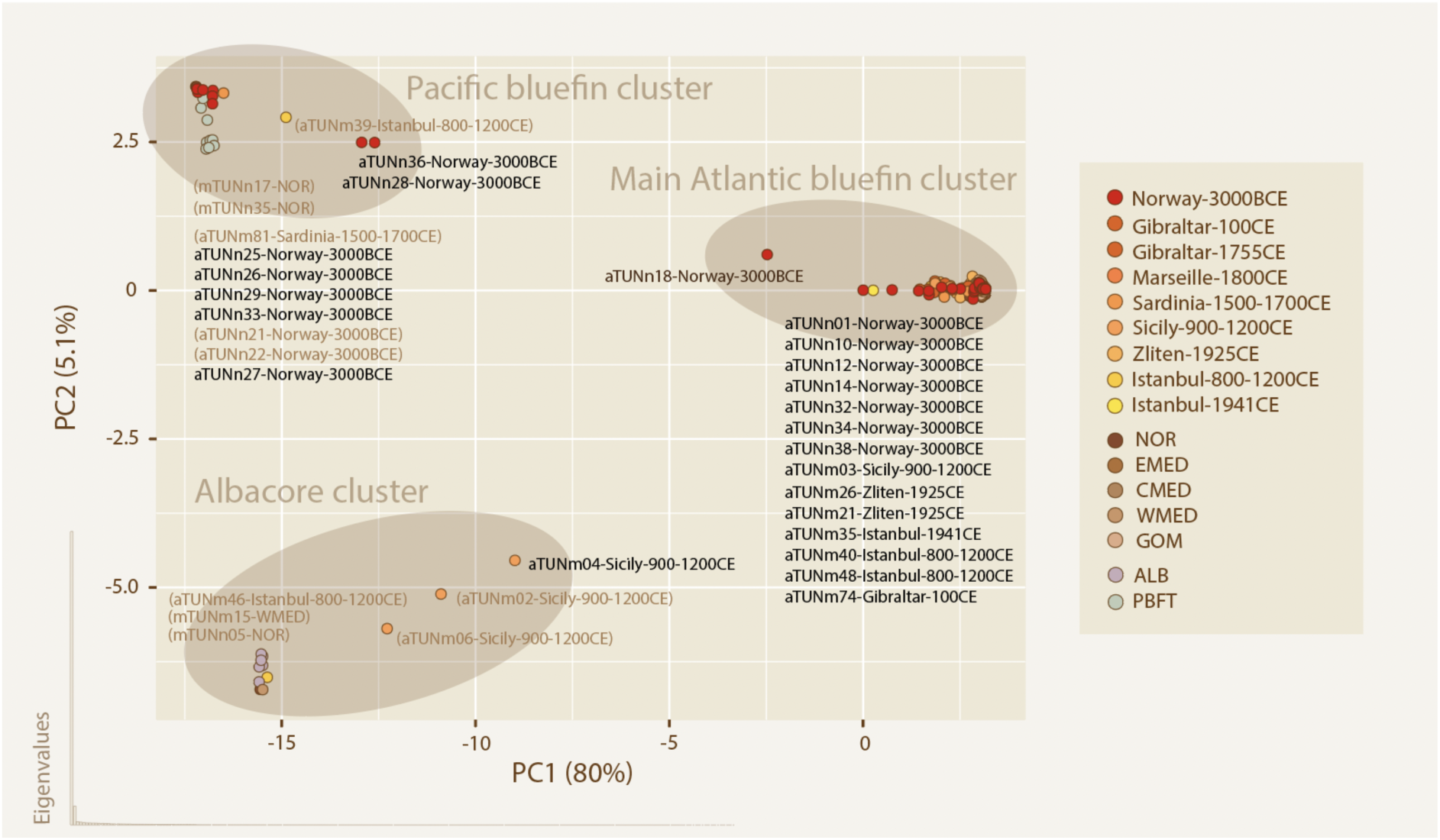
PCA of all samples included in the exploratory analysis, prior to omission of identical- and high missingness samples. The plot shows an interspecific PCA. Samples that were excluded from subsequent population genomic analyses are marked with the sample name in dark brown. Sample names in parentheses indicate diverging samples that were kept, but taken into account as introgressed haplotypes, in the population genomic analyses.

**Figure S4:**
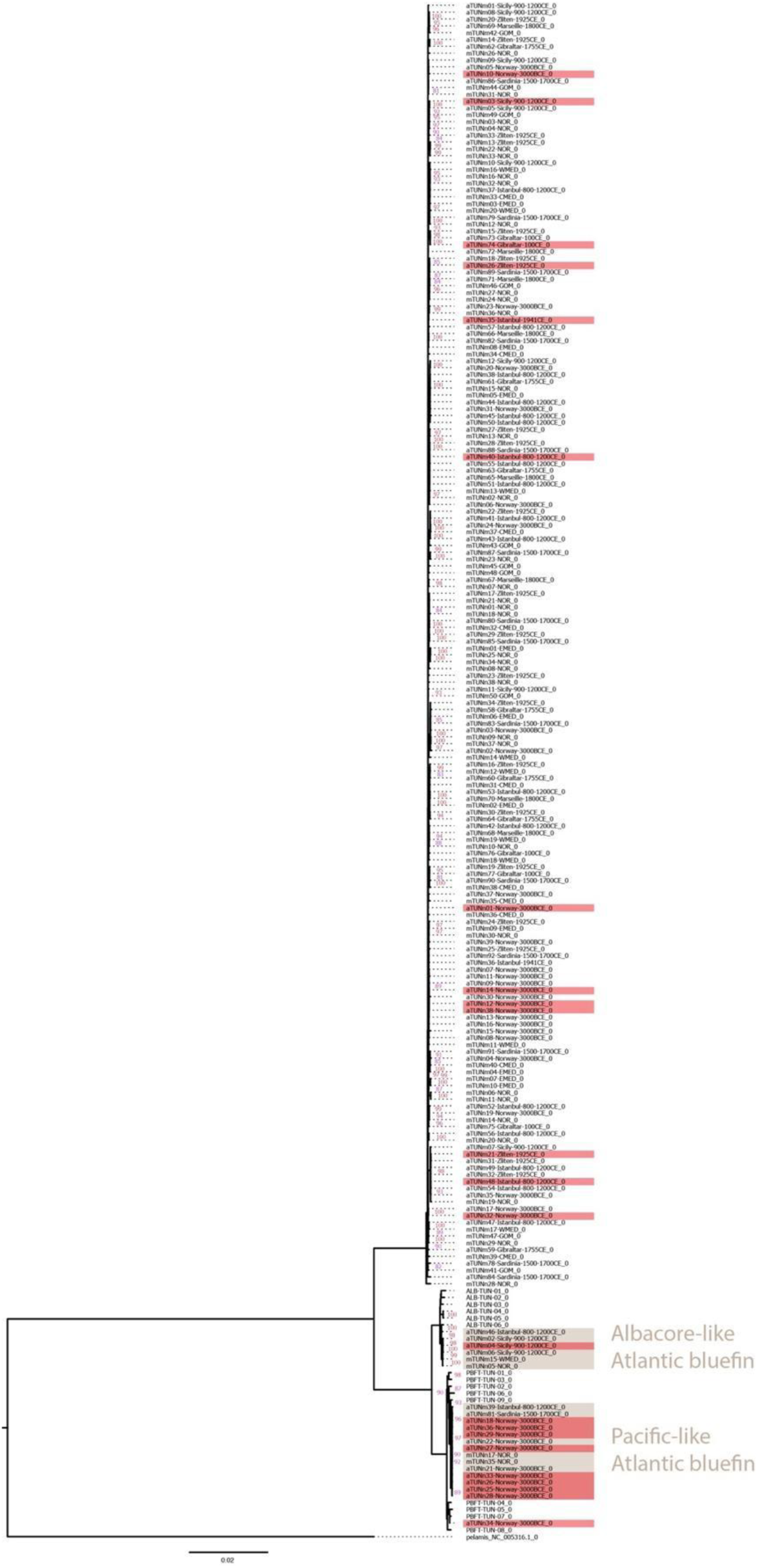
ML phylogeny of all samples included in the exploratory analysis, prior to omission of identical- and high missingness samples. Bootstrap values over 80 are shown in pink. Samples that were excluded from subsequent population genomic analyses are highlighted in red. Samples highlighted in brown indicate diverging samples that were kept, but taken into account as introgressed haplotypes, in the population genomic analyses.

**Figure S5:**
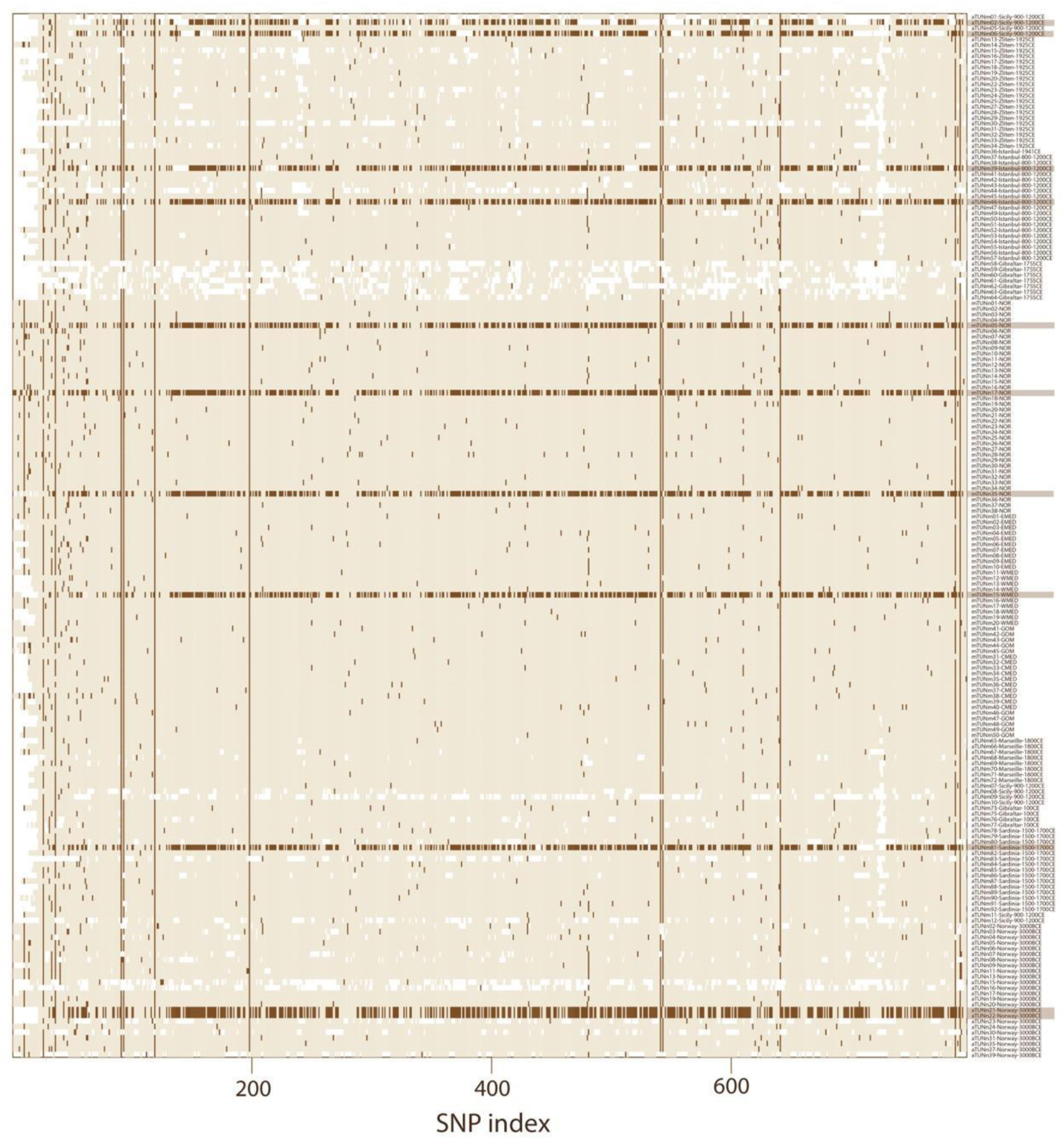
Missing loci (white) and the presence of second alleles (dark brown) across the Atlantic bluefin samples (dataset: AllABFT). The 11 specimens that were identified with introgressed MT genomes have a high number of divergent alleles when compared to the Atlantic bluefin reference genome.

**Figure S6:**
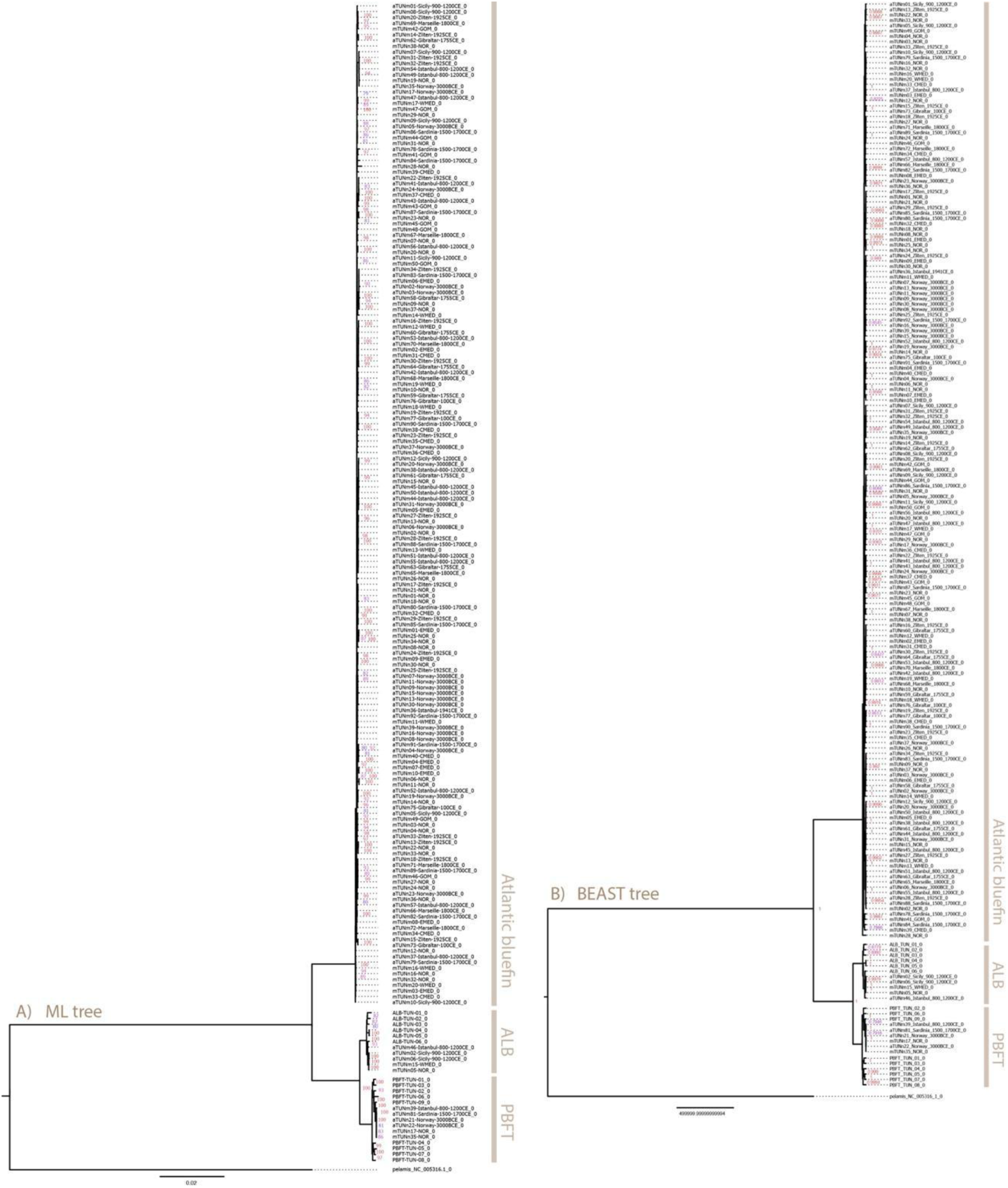
ML and Bayesian phylogenies of all samples included in population genomic analyses (dataset: All_ALB_PBFT) using Skipjack tuna (Katsuwonus pelamis) as outgroup. Bootstrap values over 80 and posterior probability values over 0.8 are shown in pink in A) and B) respectively. Species clades are highlighted in brown.

**Figure S7:**
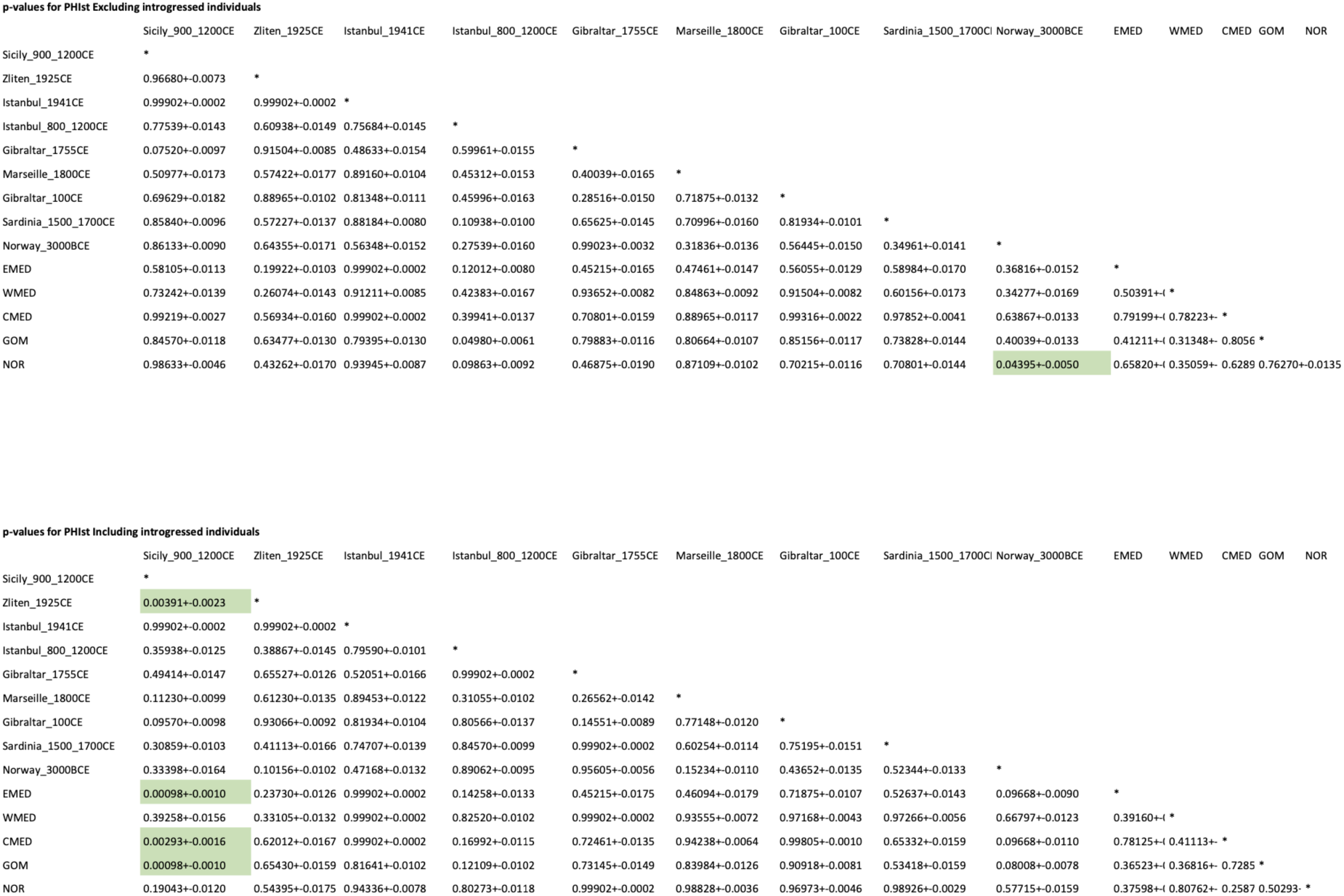
ΦST P-values from Arlequin for the AllExIntrog and AllABFT datasets, corresponding to Figure 3A) and 3B) respectively. Significant p-values are marked in green. After correcting for multiple testing (Bonniferoni correction), none of the p-values remained significant.

**Figure S8:**
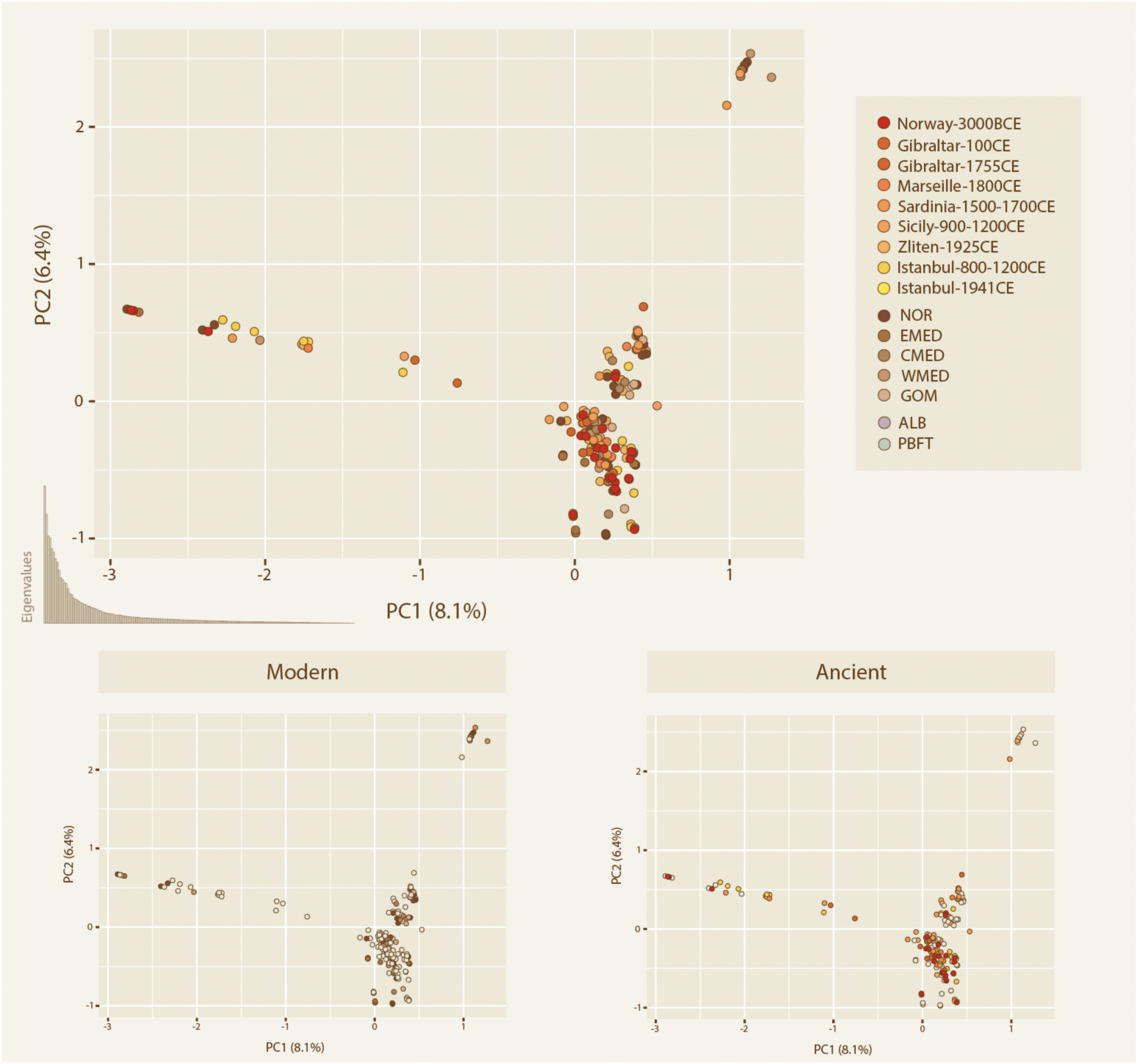
PCA of all non-introgressed Atlantic bluefin individuals (dataset: AllExIntrog). The ancient and modern samples are highlighted in the bottom panel. Eigenvalues are shown in the left corner of the upper panel.

**Figure S9:**
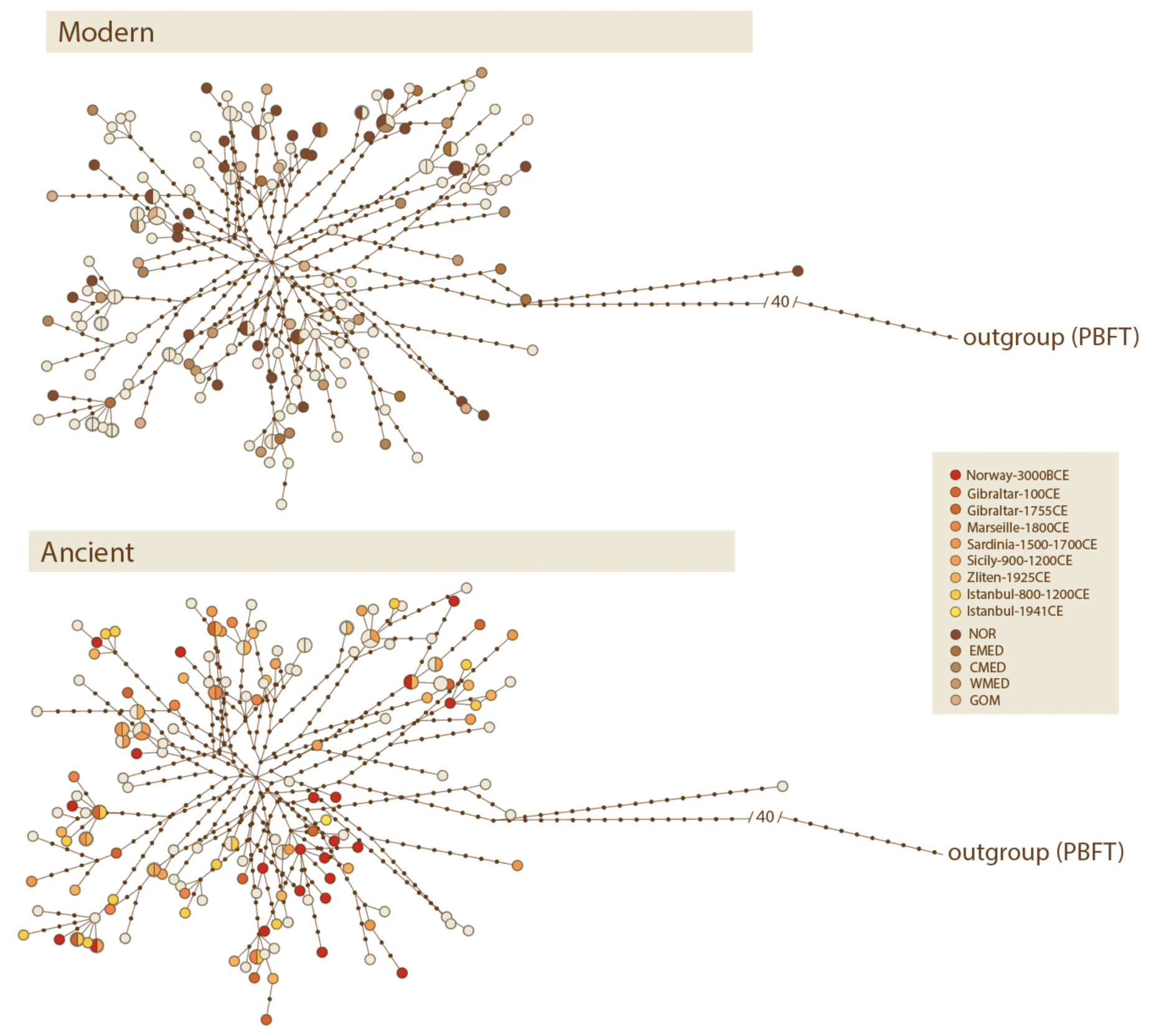
Haplotype network of all non-introgressed Atlantic bluefin individuals, using dataset AllExIntrog and Pacific bluefin (Thunnus orientalis) as outgroup. Each node represents a unique haplotype.

**Figure S10:**
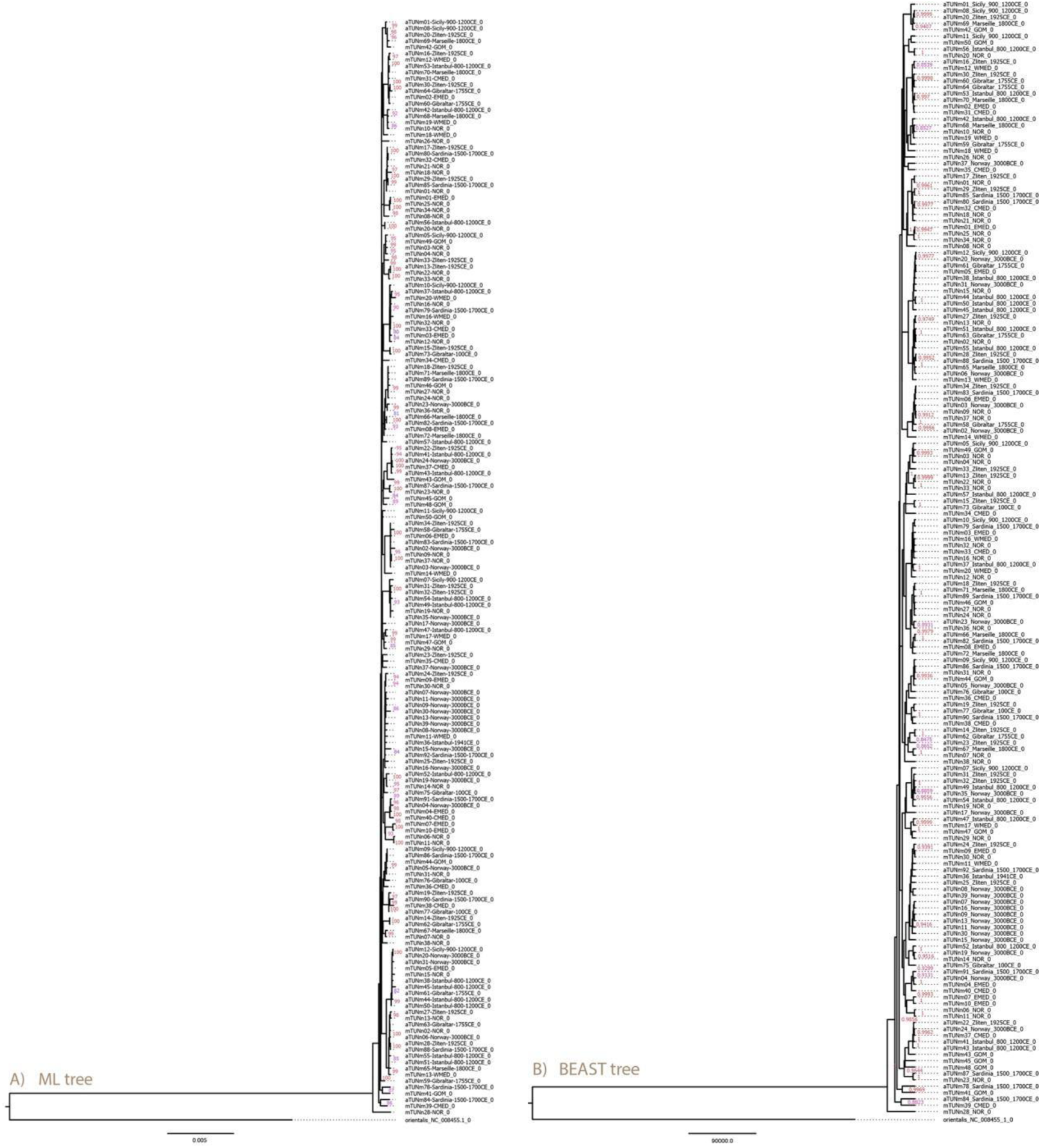
ML and Bayesian phylogenies of all samples included in population genomic analyses (dataset: AllExIntrog) using Pacific bluefin (Thunnus orientalis) as outgroup. Bootstrap values over 80 and posterior probability values over 0.8 are shown in pink in A) and B) respectively.

